# Intensity-dependent topographical expansion of sensory representations

**DOI:** 10.64898/2026.06.05.730458

**Authors:** Li-Bo Zhang, Amin Dehghani, Li Hu, Patrick Sadil, Liz Losin, Martin A. Lindquist, Tor D. Wager

## Abstract

Neuroimaging studies typically assume that sensory properties are encoded in response magnitude within fixed neural populations. However, this approach does not capture changes in the spatial extent of activation topography, despite growing evidence for its behavioral relevance. Stimulus intensity provides a powerful test case for the role of activation topography as a coding feature because it is a basic, parametrically varying property shared across sensory modalities. Using a Bayes factor-based approach and four functional magnetic resonance imaging datasets (three large-scale datasets [total N = 609] and one precision dataset [>2300 trials]), we tested whether higher-intensity stimulation is associated with expansion of activation topography. Participants received sensory stimuli of varying intensities in somatosensory (heat, laser, tactile), auditory, and visual modalities. High-versus low-intensity painful stimulation consistently produced topographical expansion in areas including the primary somatosensory, posterior midcingulate, primary visual cortices, and cerebellar lobules V and VI. This result replicated across two independent large-scale datasets and within individual participants in the precision dataset. Expansion was also observed for tactile, auditory, and visual stimulation, and its extent correlated with psychophysical discriminability. Topographical expansion involved both the enlargement of already-activated areas and the recruitment of novel regions. These findings establish topographical expansion as a replicable feature of intensity coding, challenging the prevailing assumption of a fixed neural topography.

## Introduction

Sensory stimulus properties such as intensity convey substantial information about the causes and consequences of stimulation – including whether it is safe or damaging^1^, whether its source is near or far^2^, and its communicative value^3,4^. How the brain encodes stimulus properties is thus a central question in neuroscience. The most widely studied principle is magnitude coding, in which stimulus properties are encoded by the magnitude of recorded signals^5,6^, such as single-neuron firing rates and local field potential amplitudes in invasive electrophysiological studies, and voxel activation magnitude in noninvasive neuroimaging studies. For example, the magnitude of blood-oxygenation-level-dependent (BOLD) signal in certain brain regions parametrically scales with sensory intensity^7,8^. A complementary principle, population coding, emphasizes distributed activity patterns across sets of neurons or voxels^5,6^. Consistent with this principle, multivariate neural patterns can be used to decode somatosensory^9,10^, auditory^11^, and visual stimuli^12^. However, these coding principles typically quantify response amplitude or multivariate patterns within a predefined and fixed set of neurons, voxels, or regions, leaving open whether changes in the set of responsive units (e.g., expanding activation extent in neuroimaging) also carry meaningful information despite growing evidence for the behavioral relevance of spatial features of brain activity^13,14^.

Neural representations are increasingly understood as dynamic and reorganizing across changing contexts^15–17^, challenging the notion that stimulus properties are encoded within a fixed feature space. One notable form of this dynamism is neural recruitment, whereby neural populations that were inactive become active in response to greater internal or external demands. Among all stimulus properties, intensity provides a powerful test case for neural recruitment because it is one of the most basic quantitative properties shared across all sensory modalities and permits within-task comparisons without task type confounding. Indeed, some of the earliest studies of neural coding focused on how stimulus intensity is encoded^18,19^. Neurophysiological studies have shown that high-intensity sensory stimuli can recruit additional neurons relative to low-intensity stimuli, especially in the peripheral and early sensory stages^20–26^. In neuroimaging, neural recruitment can manifest operationally as an expansion in the spatial extent of activation topography. If robust, intensity-dependent expansion of activation topography would support a more flexible coding principle complementing those based on response magnitude and multivariate patterns.

Some neuroimaging studies suggest that the extent of activation is modulated by stimulus intensity, for example, during painful^27,28^, tactile^29^, auditory^30–32^, and visual stimulation^33,34^. However, most studies have not demonstrated that larger activation extent reflects topographical expansion, as opposed to simply more robust activity in a given set of voxels, which would increase activation extent via an increase in statistical power. A stronger test of topographical expansion requires that two conditions be satisfied: (1) a given spatial unit is active in the high-intensity condition; and (2) the same unit is inactive in the low-intensity condition (i.e., evidence in favor of a null effect). Previous studies do not explicitly test the second condition and, with sample sizes typically smaller than 30, have limited statistical power to test it. These studies count statistically significant voxels in each intensity condition^28,32,33,35^. However, a non-significant p-value may reflect lack of power rather than a true absence of activation. Bayes factors (BFs) provide a principled approach to this problem because they quantify evidence for both the alternative and null hypotheses^36^.

Beyond this methodological ambiguity, topographical expansion also remains poorly characterized. Different types of expansion may exist. It could occur at the edges of regions activated by a lower-intensity stimulus, or recruit novel regions located far away from those commonly activated by both high- and low-intensity stimuli. In addition, expansion can reflect new recruitment of voxels within modality-specific regions or a broad shift across many regions. The latter is consistent with increases in diffuse neuromodulatory systems associated with arousal. For example, recent studies have demonstrated brain-wide neural activity and cerebral blood flow elicited by simple sensory stimuli^37^. Identifying these expansion subtypes may shed light on their underlying mechanisms.

To address these issues, we investigated intensity-dependent topographical expansion using a BF-based approach and four functional magnetic resonance imaging (fMRI) datasets (three large-scale datasets [total N = 609] and one precision dataset [>2300 trials from 22 participants]). Healthy participants received stimuli in multiple sensory modalities (including nociceptive, tactile, auditory, and visual) across multiple intensity levels and reported perceived ratings (Figure 1; see Table S1 for descriptive statistics of behavioral ratings). Using the BF-based approach^38^, we identified voxels that showed positive activation only in the high-intensity condition, which supports topographical expansion (Figure 1A; see Methods for details). The precision dataset further allowed us to determine whether topographical expansion generalizes across most individuals, rather than being driven by group averaging^39,40^. We used pain as the primary model system because intensity is central to pain’s function and a major target of clinical interventions^41–43^. We aimed to test three hypotheses: (1) activation topography reliably expands in high- versus low-intensity conditions for painful stimulation at the group and individual levels; (2) topographical expansion generalizes beyond pain to other modalities; and (3) expansion can be decomposed into distinct spatial types.

**Figure 1.**
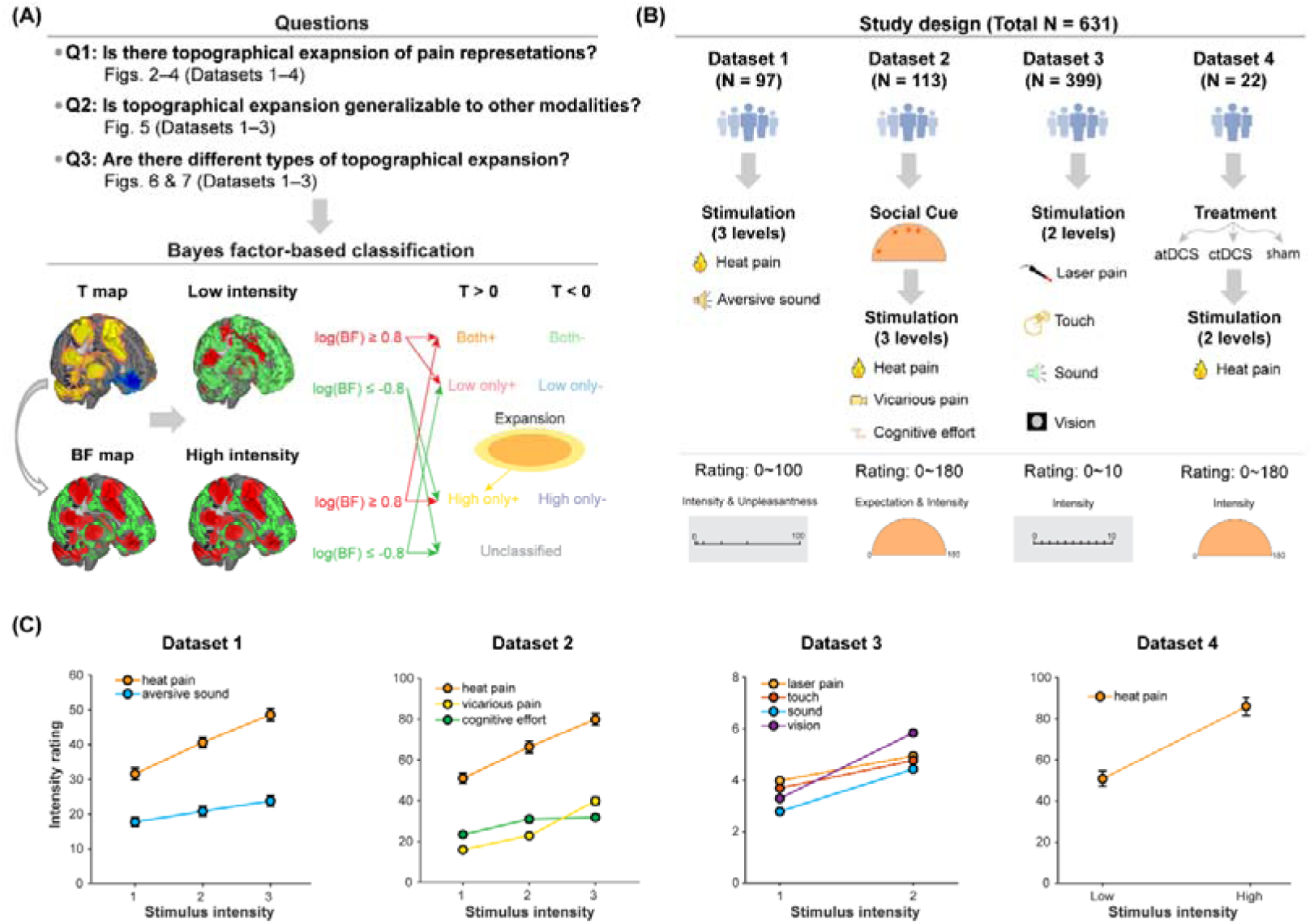
Study overview. **(A)** Research questions and analytic approach. Using four datasets, we studied whether new brain regions are recruited to process high-intensity pain stimuli compared with low-intensity pain stimuli, that is, whether brain regions activated by high-intensity pain stimuli expand topographically compared with those activated by low-intensity pain stimuli. We also examined whether it is generalizable to non-pain modalities, and whether different types of expansion can be identified. Recruitment of new regions (i.e., topographical expansion of representations) was assessed with Bayes factor-based classification of voxels. Thresholds of 0.8 and -0.8 are only examples, and slightly different thresholds were applied in different datasets (see **Methods**). **(B)** Study design of the four analyzed datasets. In all datasets, participants received painful heat stimuli of several intensity levels. Some datasets also included aversive sound (Dataset 1); vicarious pain and cognitive effort (Dataset 2) stimuli; and tactile, auditory, and visual stimuli (Dataset 3). Unlike in Dataset 1, auditory stimuli were pure tones in Dataset 3. Individualized analysis was conducted in Dataset 4, where >50 stimuli per intensity level were administered. **(C)** Intensity ratings of painful and non-painful stimuli. Error bars are standard errors of the mean. Some error bars are too small to be visually discerned (e.g., Dataset 3). atDCS: anodal transcranial direct current stimulation; ctDCS: cathodal transcranial direct current stimulation.

## Results

### Intensity-dependent topographical expansion for pain

To examine whether more intense noxious stimuli elicit topographical expansion, we applied the BF-based approach to Dataset 1^44^, in which participants (N = 97) received painful heat stimuli of three temperature levels: 47 °C, 48 °C, and 49 °C (Figure 1). The lowest intensity (47 °C) was painful for most trials in most participants, while 49 °C was very painful for virtually all participants on every trial (Figure 1C & Table S1). All three intensity levels of painful heat led to activation or deactivation in a wide range of brain regions (Figure S1). As expected^43,45,46^, activity in many brain regions correlated linearly with stimulus temperature, including regions such as primary somatosensory cortex (S1), secondary somatosensory cortex (S2), insula, anterior cingulate cortex (ACC), prefrontal cortex (PFC), thalamus, and cerebellum (thresholded with log(BF) = 0.8; Figure S1B). Most of these regions also correlated with trial-wise subjective pain ratings (Figure S1C), and the unthresholded intensity-encoding map and rating-encoding map were highly correlated (Pearson’s r = 0.86).

To demonstrate topographical expansion, we classified voxels into eight classes: (1) “High only+”, showing activation only in the high-intensity condition, where “only” means log(BF) > 0.8 (i.e., >6:1 evidence for activation) for high-intensity and log(BF) < -0.8 (i.e., >6:1 evidence in favor of null findings) for low-intensity; (2) “High only-”, showing deactivation only in the high-intensity condition; (3) “Low only+”, showing activation only in the low-intensity condition; (4) “Low only-”, showing deactivation only in the low-intensity condition; (5) “Both+”, showing activation in both conditions; (6) “Both-”, showing deactivation in both conditions; (7) “Low+ & High-”, showing activation in the low-intensity condition but deactivation in the high-intensity condition; and (8) “Low- & High+”, showing deactivation in the low-intensity condition but activation in the high-intensity condition (Figure 1A; see Methods for details). We mainly focused on “High only+” and “Both+” voxels. The former directly supports topographical expansion, while the latter reveals how newly recruited voxels relate to those active in both high- and low-intensity conditions.

In the 47 °C & 49 °C pair, 55.6% of voxels within gray matter were activated by both low- and high-intensity stimuli (“Both+”), indicating that painful stimuli are encoded mainly by a common core system regardless of their intensity (orange voxels in Figure 2A). 4.8% of voxels responded to high-intensity but not low-intensity stimuli and were thus classified as “High only+” voxels consistent with topographical expansion, including parts of the S1, dlPFC, dmPFC, superior temporal gyrus, and inferior temporal gyrus (yellow voxels in Figure 2A). This percentage was substantially greater than that expected by chance (p < 0.001, permutation test; Figure 2C). Activity in most “High only+” and “Both+” voxels also lay in the linear-intensity-effect mask (Figure S1B), indicating that high-intensity stimulation evoked larger responses in these regions than low-intensity stimulation did. Interestingly, the average activity differences between high- and low-intensity conditions in “High only+” and “Both+” regions were similar, with very large effect sizes for both (Cohen’s d = 1.29 and 1.20, respectively), suggesting that these regions are equally able to discriminate between high- and low-intensity stimuli (Figure 2D). When the intensity difference dropped from 2 °C to 1 °C, the percentage of “High only+” voxels dropped from ∼5% to ∼1% (ps = 0.057 and 0.054, permutation test; Figure S2). Other patterns of encoding were less prevalent. 1.8% of voxels, including a few regions such as the posterior cingulate cortex and precuneus, were deactivated only by low-intensity painful stimuli (light blue voxels in Figure S2A)

**Figure 2.**
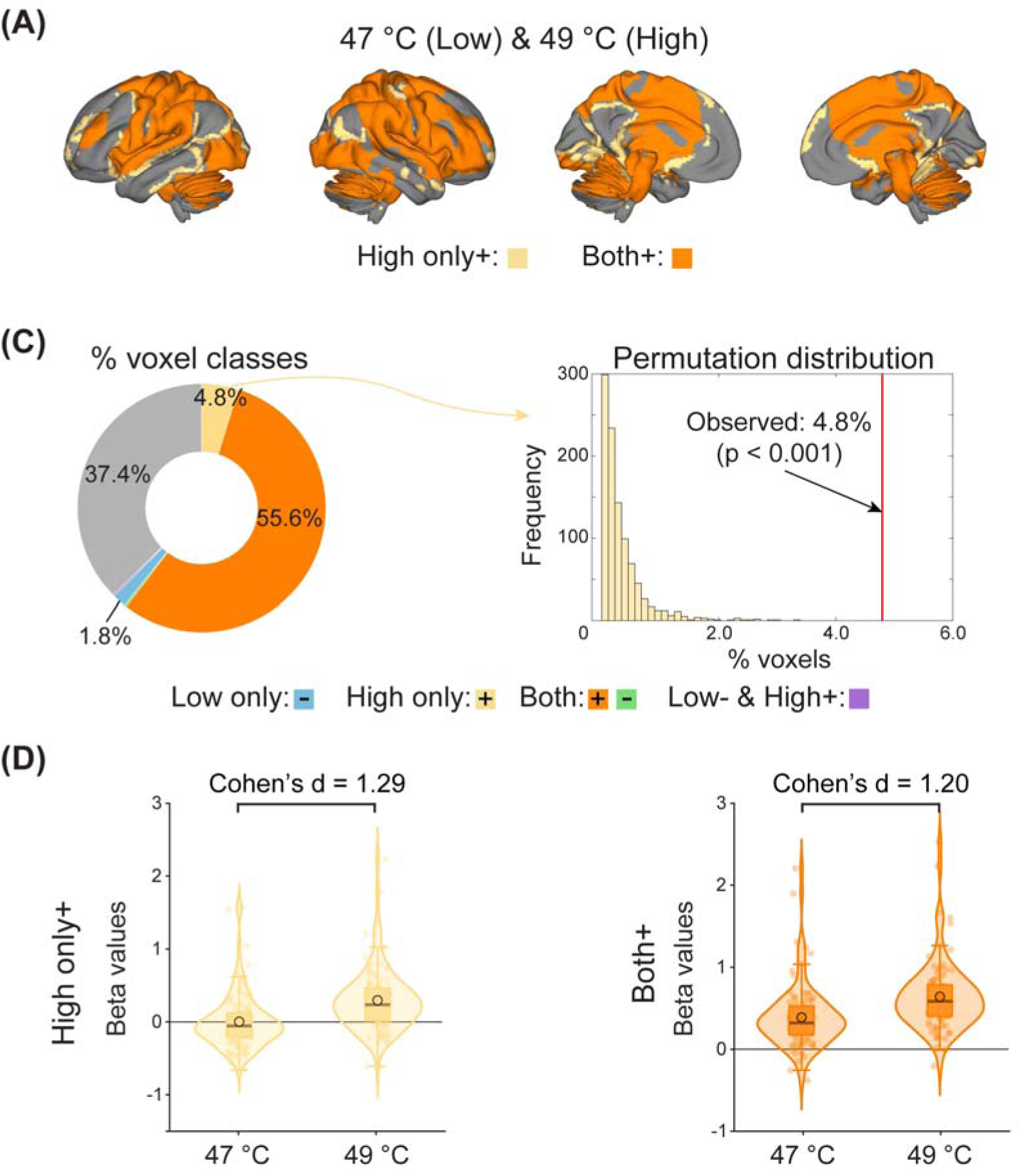
Pain-induced topographical expansion in Dataset 1. **(A)** Topographical expansion in the 47 °C and 49 °C pair. In addition to voxels (“Both+”) activated by both high- and low-intensity stimuli, some voxels (“High only+”) were activated by high-intensity stimuli but not low-intensity stimuli. Only “Both+” and “High only+” are highlighted here. See Figure S2 for all classes of voxels. **(B)** Voxel classification. 4.8% of voxels were classified as responding only to high-intensity stimuli. **(C)** Comparisons of “High only+” and “Both+” voxels. Average activity in “High only+” and “Both+” regions was similarly able to discriminate between high- and low-intensity stimuli.

Next we examined the distribution of changes across all three intensities in Dataset 1, i.e., 47 °C, 48 °C, and 49 °C. This analysis confirmed that most regions were activated at all intensities (orange in Figure S3A). However, topographical expansion occurred in different brain regions at different intensity levels. Notably, common expansions at moderate and high stimulus intensity levels were located mainly in posterior regions (medial occipital, posterior cingulate, superior cerebellum, and sensorimotor cortex; yellow in Figure S3A), whereas unique expansions with the highest-intensity stimuli occurred in dmPFC and medial temporal regions (red in Figure S3A). This suggests that distinct processes are recruited at different intensity levels rather than a single uniform gradient.

The preceding analyses operationally defined pain encoding as activation relative to implicit baselines. To evaluate the robustness of our findings, we further tested whether topographical expansion persists when employing the 47 °C condition as a baseline. Specifically, we subtracted activity elicited during this condition from the 48 °C and 49 °C conditions prior to re-running the BF-based voxel classification. We observed approximately 1.4% “High only+” voxels, notably in mPFC (Figure S3B), providing evidence that topographical expansion is related to pain processing.

To examine whether topographical expansion could be related to subjective pain perception apart from objective stimulus intensity, we trichotomized single-trial pain ratings at the 33rd and 66th percentiles and classified voxels according to their responses to low or high tertiles. The resulting voxel class map was highly similar to that of the intensity-based map (Figure S4). To further explore the topographical expansion encoding of pain perception independent of physical intensity, we median-split trial-wise pain ratings within each intensity condition and conducted BF-based voxel classification. Within each of the 47 °C and 48 °C conditions, 1.6% and 1.2% of voxels responded positively to high pain rating trials only, with evidence for null activation for low-pain stimuli at the same intensity at log(BF) < -0.8 (ps = 0.026 and 0.027, permutation test; Figure S5A-B). However, for the 49 °C condition, only 0.25% of voxels showed high-rating-only activation (p = 0.403, permutation test; Figure S5C). These results suggest that subjective pain experiences can also induce topographical expansion when the stimulus intensity is moderate, and may be saturated at very high intensities.

### Replicable topographical expansion for pain

To replicate topographical expansion across noxious stimulus intensities, we applied the same analyses to two independent datasets. In Dataset 2 (N = 113), participants were presented with one of two social cues indicating the intensity of forthcoming heat stimuli, and then received heat stimuli of three temperature levels: 48 °C, 49 °C, and 50 °C. We pooled data from both social cues to focus on the encoding of intensity. In the 48 °C & 50 °C pair, we observed similar findings to Dataset 1. As before, the majority of voxels were responsive to both high- and low-intensity heat (22.5% “Both+”; Figure 3A-B, left). 5.6% of voxels (p < 0.001, permutation test) responded only to high-intensity heat (“High only+” in Figure 3A-B, left), including voxels in S1, M1, dlPFC, early visual areas, cerebellum, and other regions. 3.6% of voxels were deactivated only by low-intensity heat (“Low only-” in Figure S6B). In the 48 °C & 49 °C and 49 °C & 50 °C pairs, we also found “High only+” voxels (Figure S6C-D), but the percentages of them (1.5% and 0.4% respectively, ps = 0.029 and 0.288, permutation test) were smaller than for the 48 °C & 50 °C pair, suggesting that the extent of topographical expansion scales with intensity differences between high- and low-intensity conditions.

**Figure 3.**
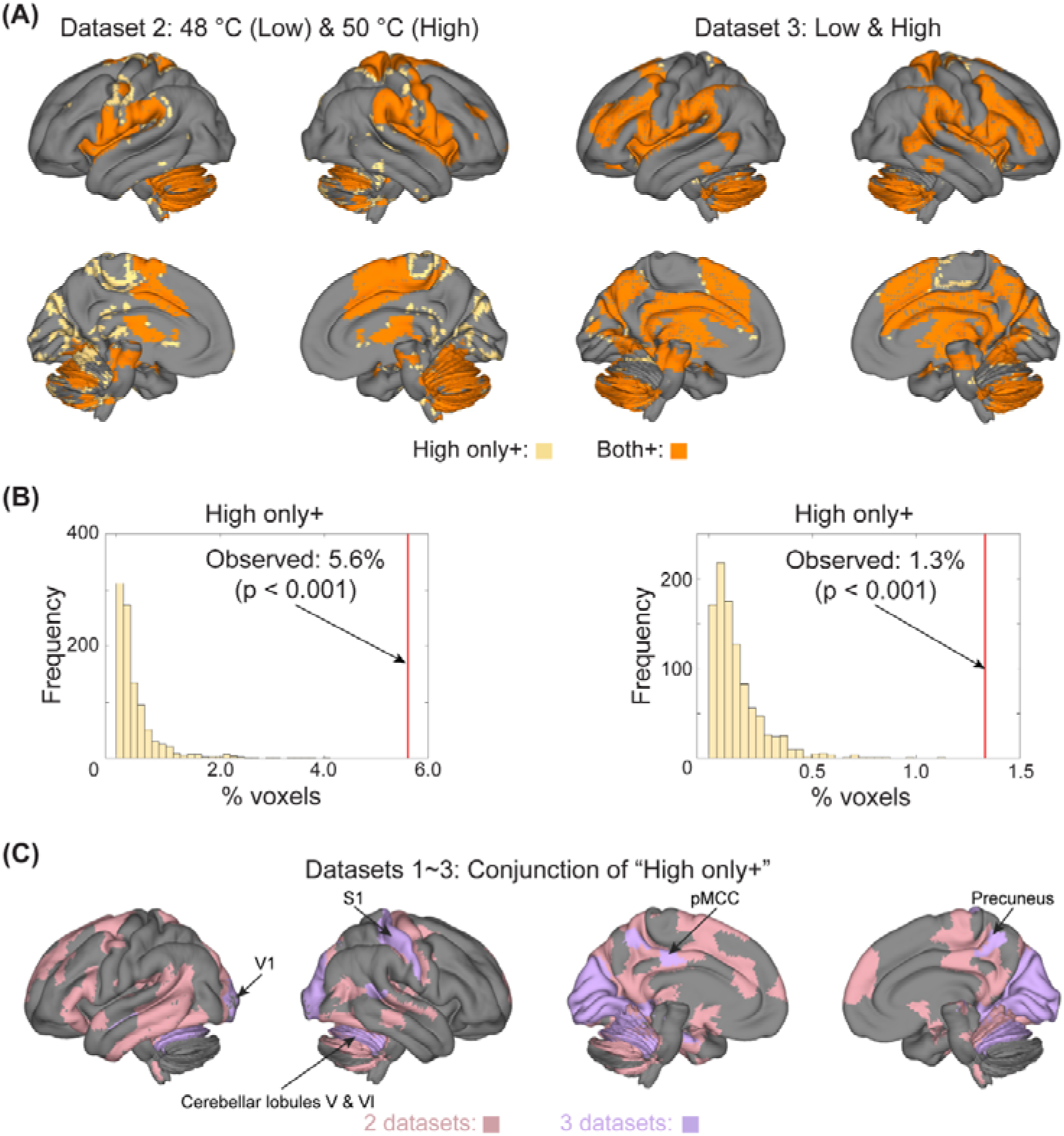
Replicable pain-induced topographical expansion in Datasets 2 and 3. **(A)** Topographical expansion in Datasets 2 and 3. In addition to voxels (“Both+”) activated by both high- and low-intensity stimuli, some voxels (“High only+”) were activated by high-intensity stimuli but not low-intensity stimuli. Only “Both+” and “High only+” are highlighted here. See Figures S6&S7 for all classes of voxels. **(B)** Percentages of “High only+” voxels. 5.6% and 1.3% of voxels were classified as responding only to high-intensity stimuli in Datasets 2 and 3, respectively. **(C)** Parcel-wise conjunction of “High only+” regions in Datasets 1–3. Regions were considered “High only+” if 5% of voxels were classified as “High only+”. Regions were masked out if they responded to high-intensity stimuli only (“High only+”) in only one dataset. pMCC: posterior mid-cingulate cortex; S1: primary somatosensory area; V1: primary visual area.

In Dataset 3, participants (N = 399) received laser heat pain stimuli of high and low intensities without additional experimental manipulations. We observed significant topographical expansion (“High only+”, Figure 3A-B, right), but with a smaller percentage (1.3%, p < 0.001, permutation test) compared with Datasets 1 and 2 (also see Figure S7 for all voxel classes). This may result from relatively smaller intensity differences in Dataset 3, as the connection between intensity differences and topographical expansion has been repeatedly observed in Datasets 1 and 2. Indeed, rating differences between high- and low-intensity pain stimuli were only 0.95 on a 0–1 scale in Dataset 3, but 17 and 28.9 in Datasets 1 and 2, which used 0–100 and 0–180 rating scales, respectively (Table S1).

The locations of “High only+” voxels differed to some degree among Datasets 1, 2, and 3 (Figures 2A and 3A), likely due to differences in samples and study design. A parcel-wise conjunction analysis identified common regions across all three datasets independently, particularly in the S1, pMCC, precuneus, V1, and cerebellar lobules V and VI (Figure 3C), suggesting that these regions were recruited uniquely by high-intensity stimuli in a generalizable fashion.

To quantitatively examine how the log(BF) threshold affects topographical expansion, we conducted a sensitivity analysis across a range of BF thresholds, from log(BF) = 0.3 – 1.2 (2x – 16x the evidence favoring the presence or absence of activation, depending on the direction). More lenient BF thresholds yielded an approximately log-linear increase in the estimated extent of expansion (Figure S8). At the conventional threshold for substantial evidence, namely log(BF) = 0.5, the amount of expansion identified was approximately double that observed with the more conservative threshold used in the main analyses.

### Painful stimulus-induced topographical expansion in individuals

The group analyses above average evidence in each voxel across individuals and are influenced by variation in inter-subject alignment. To examine expansion within individual participants, we applied the Bayesian approach to a precision fMRI pain study that included >50 trials per condition across 3 scanning days. Participants in Dataset 4 (N = 22) received individually tailored high- and low-intensity heat pain stimuli before and after sham and verum transcranial direct current stimulation (tDCS) in different sessions. Here we examine only the effect of stimulus intensity; tDCS effects will be reported in a separate paper.

We observed substantial inter-individual variability in the spatial location and amount of “High only+” voxels (Figure S9A). On average, 5.0% of voxels across gray matter in individual maps were classified as “High only+” (Figure 4A). More voxels responded to both high- and low-intensity stimuli (activation: 6.3% Both+ across gray matter, deactivation: 6.3%). Interestingly, the percentage of “High only+” voxels correlated significantly with rating differences between high- and low-intensity conditions (r = 0.58, p(FDR) = 0.03, correcting across 6 voxel classes examined), whereas the percentages of other voxel classes did not (Figure 4B; see Figure S9B for results for other voxels classes). At the group level, many areas were classified as “High only+” in more than 20% of individual participants (i.e., ≥ 5/22 participants; Figure 4C). Some of these regions, including S1, pMCC, and cerebellar lobules V and VI, were also present in Datasets 1 –3. These findings confirm the existence of topographical expansion in individualized data, suggesting that it is unlikely to be an artifact of group averaging or inter-subject misalignment.

**Figure 4.**
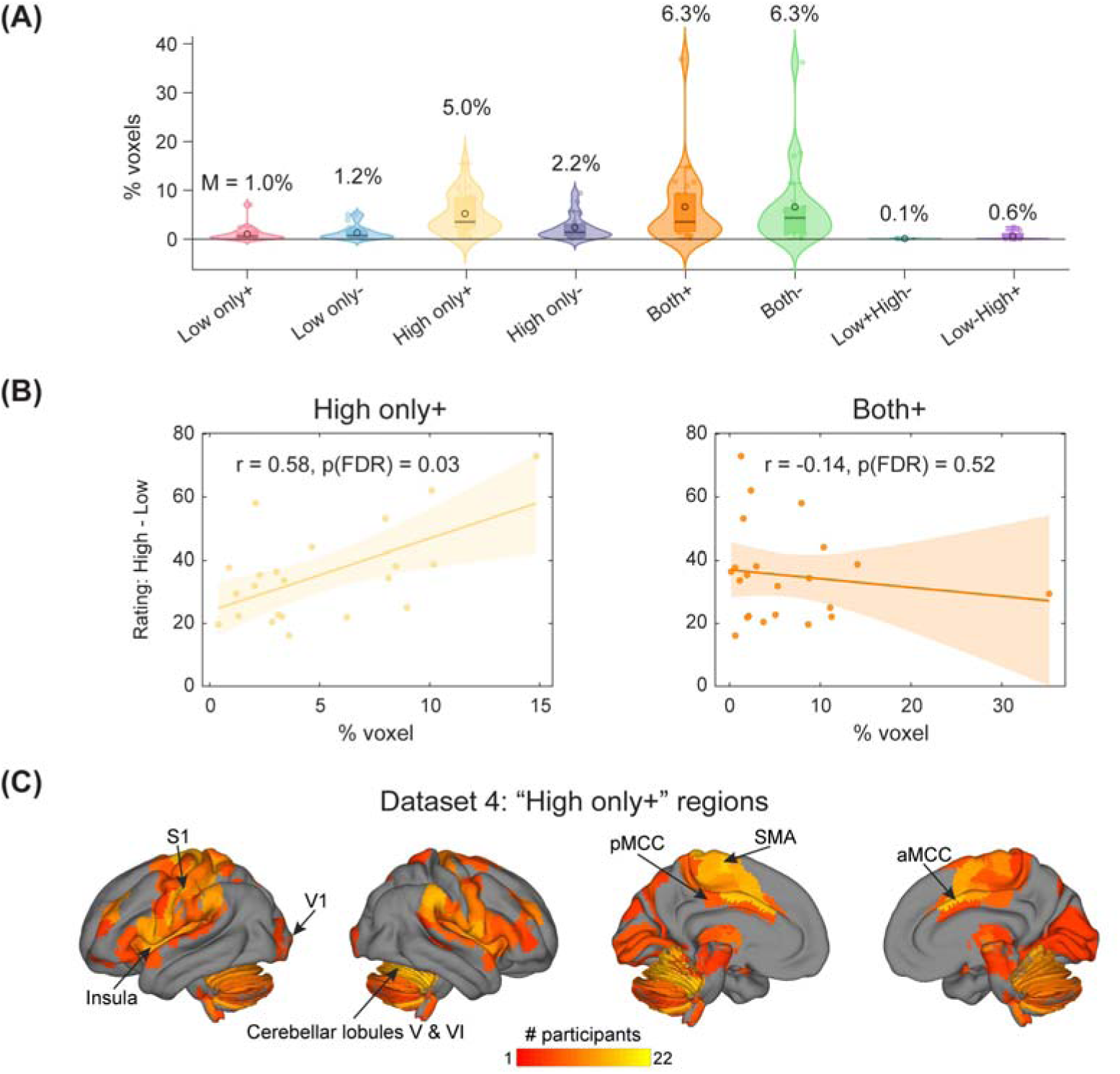
Individualized topographical expansion in Dataset 4. **(A)** Distribution of voxel classes. On average, 5.0% of voxels were activated by high-intensity stimuli only. **(B)** Correlation between “High only+”/“Both+” and rating differences. Percentages of “High only+” voxels correlated with rating differences between high- and low-intensity conditions. P values were corrected across all voxel classes except “Low+ & High-” and “Low- & High+”, as a significant number of participants (19 and 13, respectively) had no voxels within these two classes. **(C)** Number of participants showing “High only+”. Regions were considered “High only+” if 5% of voxels were classified as “High only+”. Regions were masked out if they responded to high-intensity stimuli only (“High only+”) in fewer than 20% of participants (i.e., ≤ 4 participants). aMCC: anterior mid-cingulate cortex; pMCC: posterior mid-cingulate cortex; S1 : primary somatosensory area; SMA: supplementary motor area; V1 : primary visual area.

### Potentially general topographical expansion for non-pain tasks

To examine whether topographical expansion occurs in other sensory modalities, we analyzed non-pain data in Datasets 1–3. In Dataset 1, participants received aversive auditory stimuli of three intensity levels (L1, L2, and L3). The auditory stimuli were nails scratching a chalkboard or emotionally aversive sounds (attacks, screaming, and crying) from the International Affective Digital Sounds database^47^. We pooled data from both types of auditory stimuli and classified voxels based on BFs. In contrast to pain, very few voxels (0.001%) responded uniquely to high- or low-intensity auditory stimuli, in spite of robust “Both+” activation and large stimulus intensity difference (L1 compared with L3, p = 0.998, permutation test; Figure 5).

**Figure 5.**
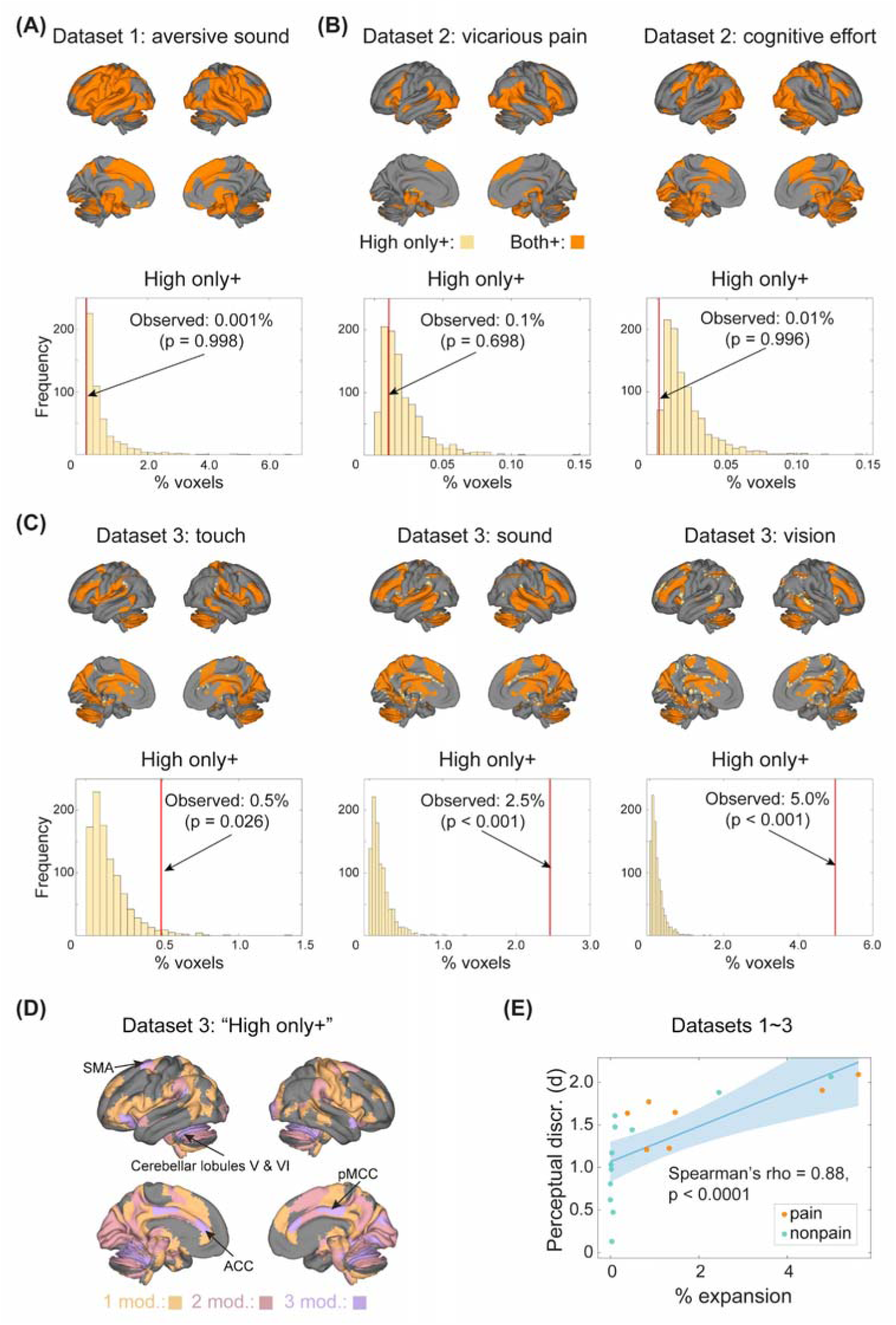
Non-pain-induced topographical expansion in Datasets 1–3. **(A&B)** Little topographical expansion for non-pain conditions in Datasets 1 and 2. Very few voxels (“High only+”) were activated by high- but not low-intensity stimuli for the aversive sound (Dataset 1), vicarious pain, and cognitive effort conditions (Dataset 2). **(C)** Topographical expansion for non-pain conditions in Dataset 3. Non-painful tactile, auditory, and visual stimuli caused topographical expansion. Unlike in Dataset 1, auditory stimuli were pure tones in Dataset 3. **(D)** Location of modality-selective and modality-general “High only+” areas. There were few “High only+” voxels across all four modalities and they were thus not visually discernible in the surface plot. No voxel was selective to audition or vision. **(E)** Correlation between perceptual discriminability and topographical expansion. Across all considered “High & Low” pairs, the extent of topological expansion correlated with perceptual discriminability, which was quantified as Cohen’s d of the rating differences between high- and low-intensity conditions. “Mod.”: modality/modalities. ACC: anterior cingulate cortex; pMCC: posterior mid-cingulate cortex; SMA: supplementary motor area.

In Dataset 2, participants watched videos of other people in pain (vicarious pain) and completed an effortful mental rotation task with three intensity levels (L1, L2, and L3, corresponding to different degrees of 3-D stimulus rotation and thus difficulty). No significant topographical expansion was found in the vicarious pain task (voxel percentage: 0.1%, p = 0.698, permutation test) and cognitive effort task (voxel percentage: 0.01%, p = 0.996, permutation test) even across large intensity differences (L1 & L3, Figure 5B-C).

However, in Dataset 3, we found topographical expansion for non-pain stimuli. In this study, participants received non-painful electro-tactile, auditory, and visual stimuli of two intensity levels each. Compared with pain (1.3% expansion; Figure 3B), more topographical expansion was observed for auditory (voxel percentage: 2.5%, p < 0.001, permutation test) and visual stimuli (voxel percentage: 5.0%, p < 0.001, permutation test). Topographical expansion was also significant but less extensive in touch (voxel percentage: 0.5%, p = 0.026, permutation test). A parcel-wise conjunction analysis revealed that regions including the ACC, pMCC, supplementary motor area (SMA), and cerebellum showed topographical expansion in three of the four modalities (nociceptive, tactile, auditory, and visual), while S1 and M1 were selectively expanded only for pain or touch (Figure 5D). These observations were confirmed by a voxel-wise conjunction analysis (Figure S10). 60.5% of “High only+” voxels were shared across at least two of the four modalities (Figure S10), with overlapping regions including the pMCC, aMCC, and SMA. No “High only+” voxels were selective to auditory or visual stimulation alone. In contrast, a number of “High only+” voxels, including those in the S1 and M1, were recruited only in pain (30.0%) and touch (9.5%).

Taken together, these findings suggest that topographical expansion occurs across sensory modalities in select cases. Topographical expansion did not seem to be a function of affective intensity, as it did not occur with highly emotionally salient affective sounds, but did occur with non-affective touch. We hypothesized that the divergent findings on topographical expansion for non-painful stimuli across studies may reflect the discriminability of high vs. low stimuli. In pain, more extensive expansion was found when the rating difference for high- vs. low-intensity stimuli was large (Figure 4B). Rating differences were small for non-pain modalities in Datasets 1 and 2, but larger for non-pain modalities in Dataset 3 (Figure 1C). To test this hypothesis, we correlated topographical expansion (% of gray-matter voxels) from all pairwise comparisons in Datasets 1-3 with the discriminability of the stimuli in ratings, quantified as Cohen’s d for high- vs. low-intensity. Expansion was strongly correlated with stimulus discriminability (Spearman’s rho = 0.88, p < 0.0001; Figure 5E), suggesting that the absence of significant expansion in Datasets 1 and 2 may be due to insufficient perceptual differentiation between intensity levels rather than a modality-specific effect.

### Two types of topographical expansion

Visual inspection indicated that there seemed to be two types of topographical expansion: (1) those that were expanded from adjacent regions and (2) those that were recruited de novo and far from regions activated at low intensity (Figures 2 and 3). To quantitatively confirm these observations, we calculated the distance between “High only+” and the nearest “Both+” voxels for pain data in Datasets 1 and 2, in which the extent of expansion was large enough (∼5% of gray-matter voxels) to meaningfully separate different types of expansion. Distances smaller than the smoothing kernel are consistent with spread from “Both+” voxels, which we term “spread expansion”. Larger distances are more consistent with the recruitment of new regions, which we term “emergent expansion”. In Dataset 1, 22.2% of “High only+” voxels were farther away from “Both+” voxels than the smoothing kernel and thus could be classified as emergent expansion, although the majority lay closer to “Both+” voxels and were thus classified as spread expansion (Figure 6A). Similar results were also obtained in Dataset 2: 32.5% of “High+” voxels were not within the smoothing kernel of the nearest “Both+” voxel (Figure 6B). The location of these two types of expansion varied across Datasets 1 and 2. In Dataset 1, we found emergent expansion in areas including dmPFC and aPFC, while in Dataset 2 we found emergent expansion primarily in visual areas. It is worth noting that we used the smoothing kernel as the criterion for distinguishing spread from emergent expansion solely for convenience. Other criteria could be adopted, provided that they preserve the intuition that “High only+” voxels located far from “Both+” voxels should not be classified as spread expansion. Under reasonable criteria, both types of expansion were arguably present in both datasets.

**Figure 6.**
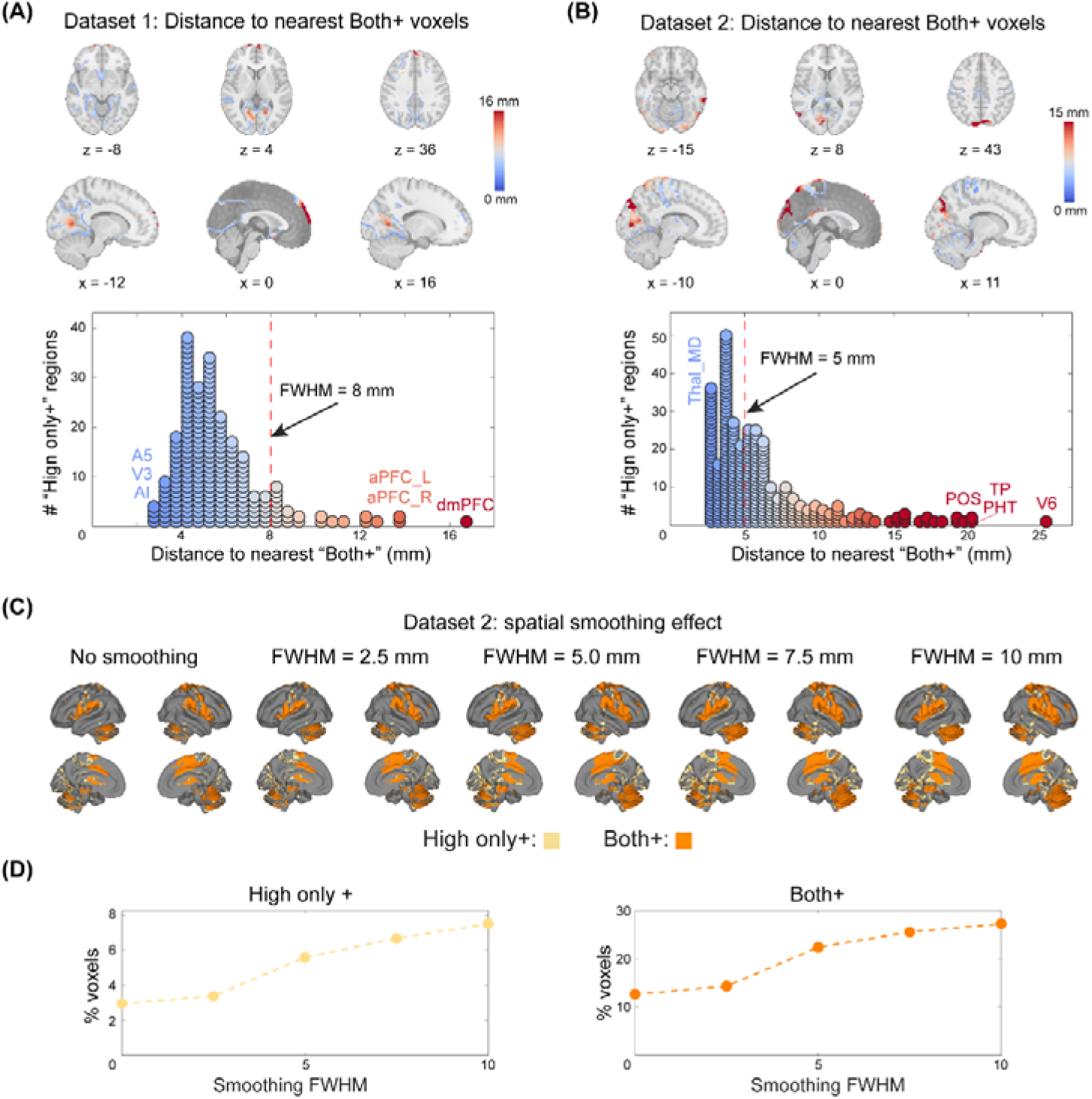
Two types of topographical expansion. **(A&B)** Distance between “High only+” and the nearest “Both+” voxels in Datasets 1 (A) and 2 (B). Some “High only+” voxels were far away from “Both+” voxels, although others were very close, suggesting distinct types of expansion. **(C)** Voxel classes under varying smoothing levels in Dataset 2. Smoothing led to the expansion of “Both+” and “High only+” regions, but some voxels were still responsive to high-intensity stimuli even when no spatial smoothing was applied. **(D)** Quantitative effect of smoothing on the percentages of “High only+” and “Both+” voxels in Dataset 2. Even without spatial smoothing, a nontrivial fraction of voxels (∼3%) responded only to high-intensity stimuli. A5: fifth auditory area; AI: anterior insula; aPFC: anterior prefrontal area; dmPFC: dorsomedial prefrontal area; PHT: posterior horizontal temporal; POS: parieto-occipital sulcus; Thal_MD: mediodorsal thalamus; TP: temporal pole; V3: third visual area; V6: sixth visual area.

The presence of emergent expansion suggests that topographical expansion reflects true neurobiological mechanisms rather than spatial blurring artifacts attributable to vascular artifacts or spatial smoothing. However, some of the spread expansion we observed may be attributable to spatial blurring. To test this possibility, we applied different smoothing kernels (0 mm, 2.5 mm, 5 mm, 7.5 mm, and 10 mm) to Dataset 2, in which unsmoothed data were readily available. Although the percentage of “High+” voxels increased as smoothing became more aggressive, the resultant voxel class maps remained similar across levels of smoothing (Figure 6C). Critically, there were spread expansion voxels even when no smoothing was applied during preprocessing. These results provide evidence that spatial smoothing alone cannot fully explain our findings regarding spread expansion.

These expansions could be driven by recruitment of isolated groups of voxels or they could reflect broader, widespread increases. Further analyses supported the latter interpretation. Figure 7A shows an oblique section through the brain for pain data in Dataset 1, illustrating activity gradients across “Both+” regions and expanded “High only+” regions. This figure reveals a broad increase in activity across regions with the higher-intensity stimulus, showing increases in voxels with evidence for null effects at low intensity as part of a smooth gradient of increases across the brain. Further, we examined the entire distribution of t-values for low-intensity vs. high-intensity stimuli (Figure 7B, left), and found that higher intensity stimuli produced a broad increase in t-values across the brain rather than an increase in a small set of voxels. The intercepts of the best linear fit between low-intensity t-values (x-axis) and high-intensity t-values (y-axis) were consistently greater than 0, indicating a brain-wide shift upwards with high-intensity stimuli, and this intercept more than doubled as the temperature difference increased from 1 °C to 2 °C (47 °C & 48 °C: y = 0.93*x + 1.57; 48 °C & 49 °C: y = 1.00*x + 1.58; 47 °C & 49 °C: y = 0.91*x + 3.24).

**Figure 7.**
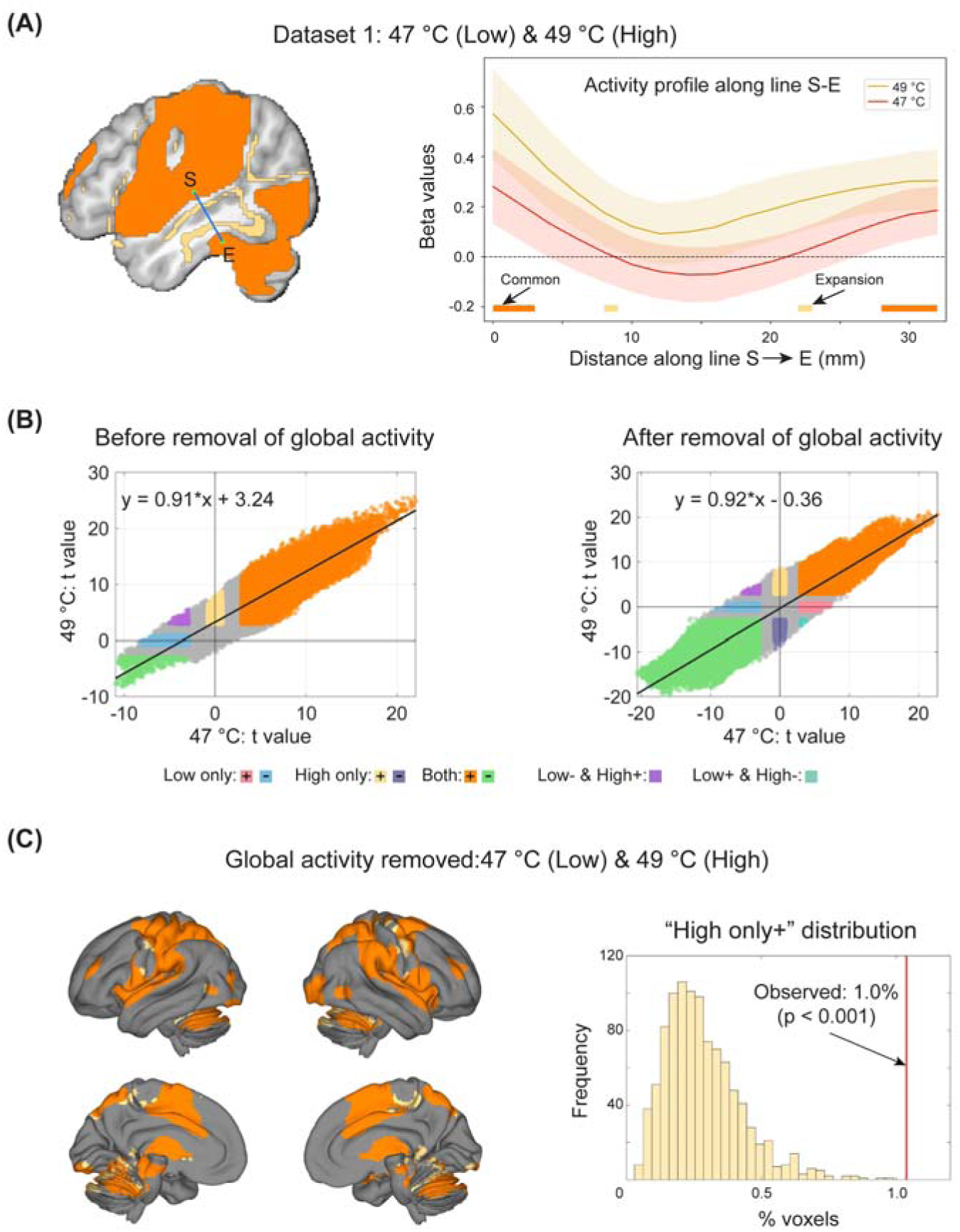
Contribution of broad activity shifts to expansion in Dataset 1. **(A)** Example activity profile across “Both+” and “High only+” voxels. Along the line connecting S (−49, −20, 7) and E (−49, −34, −22), brain activity changed systematically across “Both+” and “High only+” voxels. The 99.5% confidence intervals (CI) are plotted because they are close to the α threshold corresponding to log(BF) = 0.8. **(B)** Relationship between t value distributions in high- and low-intensity conditions before and after removal of global activity. High-intensity stimuli led to a global upward shift in brain activation compared with low-intensity stimuli. Note that the black line in the scatter plot is the fitted line. **(C)** Pain-induced topographical expansion in Dataset 1 after global activity removal. 1.0% of voxels were classified as responding only to high-intensity stimuli, even though upward shifts of brain activation were removed.

Meanwhile, the slopes remained near 1 across all three pairs of intensity, indicating that increasing stimulus intensity produced a relatively uniform increase across voxels with varying activity levels for low-intensity stimuli. Finally, when global activity within each intensity condition was removed for all participants before calculating expansion (Figure 7B, right), the extent of expansion in the 47 °C & 49 °C pair was reduced to 1% of voxels, though still significantly above chance (p < 0.001, permutation test; Figure 7C). This indicates that much – but not all – of the topographical expansion may be attributable to a brain-wide activity increase.

As in Dataset 1, expansion was also largely attributable to widespread activity increases with high-intensity stimulation for pain in Datasets 2 and 3. The percentages of voxels identified as “High only+” were reduced when global activity was removed, though some significant voxels remained (Figure S11). Similar to pain, high-intensity non-painful stimulation also produced a widespread upward shift in activity (Figure S12A-C) and removing global activity substantially reduced, but did not eliminate, topographical expansion (Figure S13). The magnitude of upward shifts also scaled with the extent of expansion before global activity removal (Spearman’s rho = 0.84, p < 0.0001; Figure S12D), further confirming the idea that a global activity increase may be partly responsible for topographical expansion. As in large-scale datasets, we also found that topographical expansion was related to global activity shifts in most individuals, and that activity increases occurred in both expanded and non-expanded areas (Figure S14). Altogether, these observations are in accord with the idea that topographical expansion is embedded in a broader activity shift across both expanded and non-expanded regions, and that broad activity shifts contribute to expansion.

## Discussion

Traditional neuroimaging analyses assume that the spatial territory over which a stimulus is represented remains largely fixed. We tested this assumption using a BF-based analytic approach across four fMRI datasets totaling 631 participants, who received high- vs. low-intensity stimulus pairs in multiple sensory modalities. We obtained four major findings: (1) the topography of pain representations expanded with increasing intensity; (2) the topographical expansion replicated in two large-scale fMRI datasets and within individuals in one precision fMRI dataset; (3) topographical expansion generalized to non-pain modalities, in proportion to the perceptual discriminability of the stimuli compared; and (4) expanded areas included the enlargement of adjacent activated areas (i.e., spread expansion) and recruitment of novel brain regions (i.e., emergent expansion). These findings establish topographical expansion as a reliable and potentially general feature of intensity representations, highlighting the dynamic nature of neural representations and challenging models that assume a fixed neural topography.

While not a central focus in traditional neuroimaging analyses, the representational value of the spatial extent of neural topography has long intrigued neuroscientists. Early work on this topic used the number of significant voxels as a measure of spatial extent^7,27,30,33,48^. However, this approach cannot distinguish true absence of activity from insufficient power to detect it, and is therefore ill-suited for identifying spatial expansion of representations. To address this limitation, we employed the BF-based approach, which can provide direct evidence for both the presence and absence of activity. With pain as the primary model system, we tested whether increasing intensity can lead to the expansion of activation topography. Across the four datasets, we found that high-intensity painful stimulation activated voxels that showed evidence for no activation (null) compared to rest with lower-intensity stimulation. This combination demonstrated recruitment of previously non-activated voxels and thus demonstrated intensity-dependent topographical expansion. The precise location of voxels showing intensity-dependent expansion varied across datasets, possibly due to the heterogeneity of study design and samples combined with the requirement of null activation at lower stimulus intensity vs. baseline. Nevertheless, at the parcel level, some areas consistently showed topographical expansion for pain. Notably, S1, pMCC, precuneus, V1, and cerebellar lobules V and VI topographically expanded across all three large-scale datasets. Additional areas, including dmPFC, dlPFC, ACC, M1, and SMA, showed expansion in two datasets. These observations were further confirmed in the precision dataset. S1, SMA, and cerebellar lobules V and VI were among the most consistently expanded areas across participants (>16 of the 22 participants).

Most of the expanded regions are relevant to pain, although their functions are not pain-specific^43,45,46^. For example, S1 is implicated in intensity processing^49–52^, the ACC is related to diverse motivated actions although it contains some pain-specific representations^53,54^, and other regions, such as the PFC, M1, SMA, and cerebellar lobules V and VI, are involved in the cognitive dimensions and sensory integration of pain^42,55–57^. V1 and other visual areas are not typically associated with pain, but some evidence suggests they may still carry information about pain^58^. Previous studies, however, have mainly associated pain with the magnitude of activation in these regions^43,59,60^. Our findings further show that the spatial extent of activation in these regions can also code pain intensity. Indeed, the extent of expansion was correlated with rating differences between high- versus low-intensity conditions, whereas no significant correlations were found for the extent of commonly activated voxels or other voxel classes. These observations indicate that topographical expansion is behaviorally meaningful, rather than a functionally inconsequential phenomenon.

Previous studies have also suggested that topographical expansion could happen in multiple domains rather than pain-specific^24,31–33^, although systematic investigations into neural recruitment across domains remain scarce. We found topographical expansion in non-pain modalities in selected cases, particularly in Dataset 3. However, not all non-pain modalities in our analyses exhibited expansion, particularly aversive sound, vicarious pain, and cognitive effort in Datasets 1 and 2. Two factors may contribute to these differential findings. First, high versus low rating differences for non-painful stimuli were smaller than those for pain stimuli in Datasets 1 and 2, but were comparable to or greater than those for non-painful stimuli in Dataset 3. Even for pain stimuli in Datasets 1 and 2, the expansion extent was smaller when intensity/rating differences were small. Across all sensory modalities, the amount of expansion was strongly correlated with rating differences (expressed in effect size, a measure of discriminability) across all modalities. Second, non-painful stimuli in Dataset 3 were simpler and better matched across intensity levels compared with those in Datasets 1 and 2. For example, auditory stimuli were pure tones of different intensities in Dataset 3, but aversive sounds of attacks, screaming, and crying in Dataset 1. Content differences in these complex stimuli may alter neural representations and overshadow intensity effects^61,62^. These findings also indicate that topographical expansion is likely to be driven by sensory intensity rather than affect, as it was most apparent for pure tones but not for affectively arousing stimuli.

Supporting the generalizability of topographical expansion, areas showing expansion partially overlapped across modalities in Dataset 3. Specifically, the SMA, ACC, PCC, and cerebellar lobules V and VI showed expanded topography in three modalities. These regions are associated with domain-general, cross-modal processes, such as the estimation of perceptual magnitude^63^, attention engagement/alertness^64,65^, and task-maintenance or control^66–69^. Such modality-general functions provide a likely mechanism for their expansion in different sensory modalities. As stimulus intensity increases, individuals allocate greater attentional and cognitive resources to external stimuli, enhancing attention to relevant features and facilitating appropriate response selection.

Similar to functional brain networks^70^, the topographically expanded areas appear to comprise two types: spread expansion and emergent expansion. The first type, including the S1 and cerebellum, was close to regions commonly activated by both high- and low-intensity stimulation; the second, including the dmPFC and some visual areas, occurred far away from the closest commonly activated areas. fMRI data are spatially smooth, and some of the apparent expansion might result from stronger activity in voxels already activated at lower intensities combined with smoothing, but sensitivity analyses indicated that smoothing effects were modest and did not produce spread or emergent expansion. In addition, spurious estimates of expansion related to spatial smoothness were mitigated by our stringent log(BF) = 0.8 threshold. Indeed, ∼10% of expansion was observed when a lenient threshold of log(BF) = 0.5 was applied.

Mechanistically, spread expansion may be caused by the recruitment of new afferent sensory neurons, such as wide-dynamic range neurons, by high-intensity stimulation^20,71,72^, as well as recruitment of cerebral neurons that only respond at a particular intensity threshold^73,74^. Evidence from somatotopic, tonotopic, and retinotopic mapping studies suggest that central sensory maps preserve neighborhood relationships in peripheral receptor populations^75–78^. High-intensity pain is perceived as spreading beyond the stimulated body area^79,80^, and newly recruited peripheral neurons may project to cortical areas neighboring those activated by low-intensity stimulation. On the other hand, emergent expansion may arise from distinct mechanisms. One plausible mechanism is the involvement of new mental processes. During high-intensity pain, the brain may allocate additional resources to cognitive regulation to manage pain or prioritize it in planning, memory formation, and decision-making. This mechanism could explain why the PFC, a region particularly important for pain regulation^81–84^, seems to be recruited anew rather than enlarged from neighboring regions. Alternatively, both types of expansion may be explained by the theory that information processing and decision-making are broadly distributed across the brain^85–87^. High-intensity stimulation may confer increased priority for processing and activate diffuse arousal systems. Consistent with this account, high-intensity stimulation was associated with a global upward shift in neural activity relative to low-intensity stimulation, and the extent of topographical expansion was markedly attenuated following the statistical removal of global activity from participant-level images.

None of these mechanisms alone is sufficient to explain topographical expansion. Recruitment of new afferent sensory neurons cannot easily account for emergent expansion, and the involvement of new mental processes do not offer a good explanation why emergent expansion areas include visual areas or why expansion can be observed for non-painful and affectively neutral sensory stimulation. Similarly, global activity changes alone fail to account for the finding that topographical expansion persisted even after controlling for global shifts. Thus, topographical expansion is likely to reflect the combined contribution of these and other mechanisms.

Taken together, our findings establish topographical expansion as a potentially general feature of intensity coding with a rigorous BF-based approach across large-scale and precision datasets. These findings suggest that topographical reorganization may be a fundamental feature of neural coding like the magnitude and pattern of activity, with potential relevance for biomarker development. Our study also has meaningful implications for spatial models in neuroimaging. Existing neuroimaging studies that incorporate spatial models primarily use them to either improve inference by modeling spatial dependence in activation maps^88–90^ or improve localization by modeling spatial variability in activation sites across individuals or studies^91–95^. Our findings reveal that spatial extent of activation topography carries representational information, underscoring the importance of follow-up studies developing new spatial models that account for topographical expansion. Furthermore, our study reveals that activation topography can shift from moment to moment as a function of stimulus intensity, highlighting the dynamic nature of neural representations. In this sense, topographical expansion represents a rapid, state-dependent form of representational change that complements the slower representational drifts observed over days to weeks^96–99^.

The present study has some limitations. First, we only examined topographical expansion in selected domains, mostly using brief sensory stimulation. While our findings suggest that topographical expansion may be a generalizable phenomenon across painful, tactile, auditory, and visual stimulation, it still needs to be tested in other cognitive and affective domains. We have shown that clear differences in the discriminability of high- and low-intensity stimuli may be a crucial driving factor for the extent of expansion. Further studies can adopt stimuli with sufficient intensity differences in other cognitive and affective domains. Second, we have focused our analyses on stimulus intensity, which is one of the most basic properties of sensory stimulation and is easily manipulated without introducing confounding changes in other sensory properties. It would be of interest to examine whether topographical expansion can be induced by other stimulus properties. Third, we only tested topographical expansion with fMRI data. It is important to note that BOLD fMRI signals only provide an indirect measure of neural activity, reflecting metabolic and hemodynamic responses rather than direct neurophysiological discharges^100^. While these measures are coupled^101,102^, a direct 1:1 mapping between BOLD signals and underlying neural activity cannot be assumed. It is therefore possible that the observed topographical expansion *does not* necessarily reflect the recruitment of novel neuronal populations, even though prior neurophysiological investigations of neural recruitment suggest that high-intensity stimulation can indeed recruit additional neurons^20,21,24^.

## Methods

### Dataset overview

We analyzed four datasets with a combined N = 631. Datasets 1, 2, and 3 have been published elsewhere and are openly accessible^44,103,104^. Dataset 4 has not been published previously. Participants in all four datasets received heat stimuli of different intensity levels and reported perceived pain intensity. Additionally, participants in Dataset 1 received aversive auditory stimuli, participants in Dataset 2 completed vicarious pain and cognitive effort tasks, and participants in Dataset 3 received electro-tactile, auditory, and visual stimuli. Note that participants did not receive painful stimuli in the vicarious pain task, but instead watched video clips of patients in pain. In Dataset 4, participants received only heat stimuli. The intensity of painful stimuli in Datasets 1, 2, and 3 was fixed for all participants: (1) Dataset 1: contact heat temperatures: 47 °C, 48 °C, 49 °C; (2) Dataset 2: 48 °C, 49 °C, 50 °C; (3) Dataset 3: laser heat energy: 3.0 J and 3.5 J, or 3.5 J and 4.0 J. By contrast, the contact heat stimulus temperature in Dataset 4 was individually calibrated using an adaptive thermal calibration procedure, with stimulation levels selected based on participant-specific pain and tolerance thresholds and corresponding predicted pain intensity estimates.

### Participants

Dataset 1 comprised 97 individuals (47 males, age 19–54 years; Mean ± SD hereafter = 28.98 ± 5.56 years), including 33 self-reported African Americans (15 males), 32 non-Hispanic White Americans (16 males) and 32 Hispanic Americans (16 males). Participants were recruited from the greater Denver area. None of them reported neurological or psychiatric diagnoses in the past 6 months, and all reported not currently using psychoactive or pain medications. They also reported no pain-related medical conditions, no reason to believe they would be especially sensitive or insensitive to contact heat, and no current unusual pain. Eight and six participants were excluded from the main analyses due to insufficient trials (< 3) for pain and sound conditions, respectively, after removing trials with high variance inflation factor (VIF > 2.5). This left 89 and 91 participants for the pain and sound conditions.

The University of Colorado Boulder Institutional Review Board approved the study, and written informed consent was obtained from all participants.

Dataset 2 included 113 participants (44 males, 68 females, 1 other; age: 24.7 ± 5.5 years). Participants were recruited from the Dartmouth College student body and surrounding communities in New Hampshire and Vermont. Participants were healthy individuals with normal or corrected-to-normal vision and hearing, no psychiatric or neurological diagnoses within the past six months, no MRI contraindications, and no chronic pain. The Institutional Review Board of Dartmouth College approved the study, and all participants provided written consent.

Dataset 3 included 399 participants. Two participants did not provide the necessary demographic information. Of the remaining participants, 238 were females, with a mean age of 21.3 ± 3.8 years. Participants were recruited from Dalian, Liaoning Province, China. All participants were healthy and free of pain, neurological, or psychiatric disorders. The Ethics Committee of the Institute of Psychology, Chinese Academy of Sciences approved this study, and all participants gave written informed consent prior to the experiments.

Dataset 4 included 22 participants (13 males, 9 females, age: 23.8 ± 2.7 years). Participants were recruited from Dartmouth College. Participants were healthy individuals with normal or corrected-to-normal vision and hearing, no psychiatric or neurological diagnoses within the past six months, no MRI contraindications, and no chronic pain. The Institutional Review Board of Dartmouth College approved the study, and all participants provided written consent.

### Sensory stimulation and tasks

Painful heat stimuli were used in Datasets 1–4. Datasets 1, 2, and 3 also included non-painful components: (1) Dataset 1: aversive auditory stimuli; (2) Dataset 2: vicarious pain tasks and cognitive effort tasks; (3) Dataset 3: electro-tactile, auditory, and visual stimuli.

In Dataset 1, heat stimulation was delivered to four evenly spaced locations (one per run) on the volar surface of the left forearm using a 16 × 16 mm^2^ (model ATS) contact Peltier thermode (Medoc, Inc.). The temperatures were fixed for all participants, viz., 47 °C, 48 °C, and 49 °C. All heat stimuli included a plateau at the target temperature flanked by 1.7 s ramp periods with a 32 °C baseline. Heat stimuli were delivered with three different temporal profiles: (1) short: 8 s, 4.6 s plateau; (2) long: 11 s, 7.3 s plateau; (3) offset: 11 s, 7.3 s plateau, with a 1 s, 1 °C temperature spike. We disregarded these temporal differences and considered only the temperature of heat stimuli in our analyses. Each heat trial was preceded by a cue signaling the onset of heat stimuli, and all parts of the trial were separated by variable delays to allow for effective deconvolution of the BOLD signal associated with each trial element. Two kinds of aversive auditory stimuli were also administered for 8 s. The first kind was a physically aversive recording of nails scratching a chalkboard, which was derived from a study of the psychoacoustics of aversive sounds and played at three different levels of intensity in steps of 5 dB^105^. The second kind of auditory stimulus was a subset of emotionally aversive sounds (attacks, screaming, and crying) from the International Affective Digital Sounds database that had the highest arousal and lowest pleasure^47^. The arousal–pleasure difference scores were used to determine aversive sound intensity levels.

In Dataset 2, heat stimuli were administered to the glabrous skin of the ventral surface of the left forearm using a TSA2 system (Medoc) with a 16-mm Peltier contact thermode. The temperatures of stimuli were fixed at 48 °C, 49 °C, and 50 °C, with the baseline temperature at 32 °C. These stimuli lasted for 9 seconds with a 5-second plateau. To account for the delay in reaching the intended temperature, an additional two seconds were padded to both the ramp-up and ramp-down phases to ensure that heat stimuli were 9 seconds long with 5 seconds of peak intended temperature. In the vicarious pain task, participants watched video clips of patients in pain, selected from the UNBC-McMaster shoulder pain expression archive database and categorized into three stimulus intensity levels using the pre-normed self-reported pain rating and observer-estimated pain rating provided in the dataset^106^. In the cognitive effort task, images for the mental rotation task were selected from the Ganis & Kievit dataset^107^. The three intensity levels were set to 50°, 100°, and 150° of rotation.

In Dataset 3, laser heat pain stimuli were transient radiant heat pulses (wavelength: 1.34 μm; pulse duration: 4 ms) generated by an infrared neodymium yttrium aluminum perovskite (Nd:YAP) laser (Electronic Engineering, Italy). The laser beam was transmitted by an optic fiber, and its diameter was set at approximately 7 mm. Laser pulses were delivered to a predefined square (5×5 cm^2^) on the left-hand dorsum. After each stimulus, the laser beam was displaced by approximately 1 cm in a random direction to avoid nociceptor fatigue or sensitization. Two stimulus energies (3.0 J and 3.5 J) were used in 212 participants as they were relatively more sensitive to laser pain and the energy level of 4.0 J was avoided. Two other stimulus energies (3.5 J and 4.0 J) were used in the remaining 187 participants as they were relatively less sensitive to laser pain. Laser heat and contact heat stimuli both evoke heat pain, but differ in two main aspects. First, laser stimuli can selectively activate nociceptors without eliciting tactile sensations, while contact heat stimuli inevitably activate the touch receptors^108^. Consequently, brain activity evoked by contact heat could be less specific to nociception compared with that evoked by laser heat. Second, laser stimuli used in Dataset 3 were extremely brief (4 ms), while contact heat stimuli in other datasets lasted for several seconds. Non-painful stimuli were also delivered. Non-nociceptive tactile stimuli were constant-current, square-wave electrical pulses (duration: 1 ms; model DS7A, Digitimer, UK) delivered through a pair of skin electrodes (1 cm interelectrode distance) placed on the left wrist, over the superficial radial nerve. The same two stimulus intensities (2.0 mA and 4.0 mA) were used in both participant groups. Auditory stimuli were brief pure tones (frequency: 800 Hz; duration: 50 ms; 5-ms rise and fall time) delivered through headphones. The same two stimulus intensities (76 dB SPL and 88 dB SPL) were used for all participants in both participant groups. Visual stimuli were brief flashes of a gray round disk on a black background (duration: 100 ms) on a computer screen. The stimulus intensities were adjusted using the grayscale of the round disk, corresponding to RGB values of (100, 100, 100) and (200, 200, 200), respectively, for all participants in both participant groups. Stimulus intensities of tactile, auditory, and visual stimuli were determined based on a pilot behavioral experiment to ensure that the perceived ratings of low- and high-intensity stimuli were approximately 4 and 7 out of 10, respectively.

In Dataset 4, heat stimuli were administered to the glabrous skin of the ventral surface of the right forearm using a TSA2 system (Medoc) with a 16-mm Peltier contact thermode. Stimulus temperatures were individually calibrated for each participant based on pain threshold and tolerance assessments conducted prior to the experimental sessions. Thermal stimulation was delivered to three distinct skin sites on the right forearm, with site locations selected during calibration to minimize habituation and sensitization effects. Each thermal stimulus lasted 13 seconds, consisting of a 10-second plateau at the target temperature and 1.5-second ramp-up and ramp-down phases.

### Rating scale

Participants were asked to rate the perceived stimulus intensity (Datasets 1, 2, and 4) and unpleasantness (Dataset 1) with a generalized labeled magnitude scale (gLMS), which appears to be a ratio scale^109^. In Dataset 1, the scale ranged from 0 to 100, with 0 meaning “No experience”, 1.4 “Barely detectable”, 6 “Weak”, 17 “Moderate”, 35 “Strong”, 53 “Very strong”, and 100 “Strongest imaginable experience”. In Datasets 2 and 4, the scale was surrounded by a semi-circle and ranged from 0 to 180, with 0 representing “No sensation”, 3 “Barely detectable”, 10 “Weak”, 29 “Moderate”, 64 “Strong”, 98 “Very strong”, and 180 “Strongest sensation of any kind”. In Dataset 3, the 0–10 Numeric Rating Scale was used, where 0 meant ‘‘no sensation’’, and 10 meant ‘‘the strongest sensation imaginable’’.

### Study design

Detailed study designs for Datasets 1, 2, and 3 were reported in previous studies^44,103,104^. Here we briefly describe the study designs (Figure 1B); only the study design of Dataset 4 will be detailed.

In Dataset 1, participants received a total of 36 heat trials and 24 auditory trials in four runs^44^. In each run, participants received one trial at each temperature (3 temperatures in total) with each temporal profile (3 temporal profiles in total) and each kind of auditory stimulus (2 kinds in total) at each intensity (3 intensity levels in total). In other words, each run consisted of 9 heat stimuli and 6 auditory stimuli. The heat stimulation site changed for each run. Trial order was randomized, following the predetermined constraint that each temporal profile of heat stimuli and each kind of auditory stimulus were evenly split between and randomly distributed within the first and second halves (for Short and Long temporal profiles) or between thirds (for Offset temporal profile) of each run. At the very beginning of each run, a single 49 °C stimulus was delivered for 11 s to allow for the initial habituation of the skin site to contact heat. This ‘washout’ stimulus was left out of all analyses.

In Dataset 2, participants completed three tasks in three separate sessions: somatic pain, vicarious pain, and cognitive effort, each with three intensity levels^103^. Each task consisted of 72 trials in total (2 cues x 3 stimulus intensities x 2 repeats per run x 2 runs x 3 sessions). In each trial, participants first viewed a low- or high-intensity social cue for 1 s indicating the upcoming stimulus (somatic pain, vicarious pain, or cognitive effort [i.e., mental rotation task]), then reported expected intensity ratings in a 4-s expectation period, received the corresponding stimulus for 9 s, and lastly gave perceived intensity ratings for 4 s. This dataset came from a larger project that included several additional components. These data were not analyzed as they were irrelevant to the present study.

In Dataset 3, participants received 80 transient stimuli of four different sensory modalities (nociceptive laser, non-nociceptive tactile, auditory, and visual) divided into two runs, and then rated their perceived intensity with a rating scale ranging from 0 (‘‘no sensation’’) to 10 (‘‘the strongest sensation imaginable [in each stimulus modality]’’)^104^. For each sensory modality, two stimulus intensities (i.e., high and low) were delivered. In other words, each participant underwent eight experimental conditions (4 modalities × 2 intensities), with 10 trials per condition.

In Dataset 4, participants completed pain testing across three separate experimental sessions following an initial calibration session. In each session, participants underwent transcranial direct current stimulation (tDCS) targeting the left primary motor cortex (M1; C3–Fp2 montage, 2 mA) for 20 minutes. The stimulation condition (anodal, cathodal, or sham) was randomized across sessions. In the sham condition, stimulation consisted of a brief ramp-up and ramp-down period (30 s) with approximately 1 minute of active stimulation. Following tDCS, participants were positioned in the MRI scanner and completed three pain runs. Each run consisted of 12 trials administered to the right forearm, including 6 low-temperature and 6 high-temperature trials, with temperatures individually determined based on each participant’s pain threshold and tolerance obtained during the calibration session. Each trial lasted 40 s and began with a warning cue (“Get Ready”) presented for 1 s, followed by thermal stimulation lasting 13 s (including a 1.5-s ramp-up, 10-s plateau at target temperature, and 1.5-s ramp-down). This was followed by a pain rating period lasting 6 s, during which participants responded using their left hand. Across each session, participants completed 36 trials in total (18 low- and 18 high-temperature trials). During all three sessions, each participant completed 108 trials (54 low-temperature and 54 high-temperature trials). These data were drawn from a larger project that included additional experimental components (including pre-stimulation pain assessments) conducted in the same sessions; however, only post-stimulation fMRI runs were included in the present analyses.

### MRI Acquisition

Dataset 1 was collected on a 3T Siemens Trio MRI scanner at the University of Colorado Boulder (voxel size in functional images: 3 × 3 × 3 mm³ for the first 25 participants and 3.4 × 3.4 × 3.4 mm³ for the remaining participants). Dataset 2 was collected on a 3T Siemens MAGNETOM Prisma MRI scanner at Dartmouth College (voxel size in functional images: 2.7 × 2.7 × 2.7 mm³).

Dataset 3 was collected on a GE MR 750 at the Research Center of Brain Cognitive Neuroscience at Liaoning Normal University, China (voxel size in functional images: 3.0 × 3.0 × 3.0 mm³). Dataset 4 was acquired on a 3T Siemens MAGNETOM Prisma MRI scanner with 32-channel parallel imaging at the Dartmouth Brain Imaging Center (voxel size in functional images: 2.7 × 2.7 × 2.7 mm³). Details of the MRI acquisition parameters can be found in Tables S2 and S3.

### Image preprocessing

Preprocessed data were analyzed for Datasets 1, 2, and 3. Note that we did rerun preprocessing with the same pipeline. This decision introduced pipeline heterogeneity, which strengthens the pipeline generalizability of our findings, as we did not pool the data but analyzed each dataset separately^110^.

Brain images in Dataset 1 were preprocessed with Statistical Parametric Mapping 8 (SPM8; Wellcome Trust Centre for Neuroimaging, London). The preprocessing steps included: (1) removing initial images (20 images for the first 25 participants; 7 images for the rest); (2) slice-timing correction with the first image (not for the first 25 participants who were scanned with the multi-band sequence); (3) co-registration with an iterative mutual information algorithm with manual adjustment of the registration starting point; (4) normalization to the MNI-152 template (resampling voxel size = 2 × 2 × 2 mm^3^); (5) smoothing with an 8 mm full-width at half maximum (FWHM) Gaussian kernel. Image intensity outliers were identified using the Mahalanobis distance computed from a matrix containing the slice-wise mean intensity and standard deviation.

For Dataset 2, preprocessing was conducted with fMRIPrep ver 25.1.4^111^. The preprocessing included: (1) motion correction with MCFLIRT (FSL ver 7.4.1); (2) co-registering to the T1 reference using the boundary-based registration method (BBR) (FreeSurfer); (3) normalizing to the standard space; (4) smoothing with a 5 mm FWHM Gaussian kernel. Note that we also smoothed images with 0 mm (i.e., no smoothing), 2.5 mm, 7.5 mm, and 10 mm

Gaussian kernels to examine the effect of spatial smoothing on results from Dataset 2. For Dataset 3, fMRI data were preprocessed using Statistical Parametric Mapping 12 (SPM12). The preprocessing included: (1) removing the first three volumes in each run; (2) slice-timing correction using the second slice and realignment to the mean slice; (3) co-registration; (4) normalizing to the Montreal Neurological Institute (MNI) space (resampling voxel size = 3 × 3 × 3 mm^3^); (5) regressing out five principal components of the white matter (WM) and cerebrospinal fluid (CSF) signals, and the six motion parameters; (6) smoothing with a 6 mm full-width at half maximum (FWHM) Gaussian kernel.

For Dataset 4, we discarded the first ten images, along with those automatically removed by the MRI scanner, to ensure the signal reached a steady state. Then, following fMRIPrep ver 23.1.3 preprocessing, multi-echo fMRI data were further denoised using TE-dependent multi-echo independent component analysis (ICA) implemented in tedana (ver 23.0.1). To distinguish BOLD-related (T2*-dependent) signal fluctuations from non-BOLD (S0-dependent) noise, principal component analysis was first performed using the Minimum Description Length (MDL) criterion to determine the optimal number of components, followed by independent component analysis decomposition and automatic component classification by tedana. Components identified as noise were removed, and the remaining components were recombined to produce an optimally denoised time series. The resulting denoised data were then spatially normalized to MNI space using antsApplyTransforms (ANTs) and transformations derived from fMRIPrep.

### fMRI data analysis

#### First-level analysis

We ran single-trial analyses for Datasets 1 and 2, and standard first-level analyses for Datasets 3 and 4. Least Squares-All (LSA) was employed to estimate single-trial fMRI responses for Dataset 1, and Least Squares-Separate (LSS) was used for single-trial analyses for Dataset 2^112^. In LSA, trial-specific regressors for all trials were included in a single design matrix, whereas in LSS, a separate general linear model (GLM) was estimated for each trial with the target trial as a regressor of interest and all other trials collapsed into another regressor. This heterogeneity, while hampering comparability to some extent, improved the analytic generalizability of our findings^110^.

The modeling was performed with SPM8 for Dataset 1. We set a boxcar regressor for the duration of the pain or non-painful stimulation period for each trial, which was convolved with the canonical hemodynamic response function. Nuisance regressors included: (1) the six motion parameters, their mean-centered squares, their derivatives, and their squared derivatives; (2) indicator regressors for signal intensity outliers; (3) run regressors representing each run; (4) regressors modeling linear drift across the duration of each run; and (5) irrelevant task components such as the cue preceding stimulation, the first “washout” heat stimulus, the rating period, and the screen indicating the end of the task in Dataset 1. A high-pass filter of 244 s (Dataset 1) or 180 s (Dataset 4) was also applied. Estimated single-trial responses were removed from further analyses if the variance inflation factor (VIF) was greater than 2.5.

LSS modeling was conducted with nilearn ver 0.11.1 for Dataset 2. We set boxcar regressors for pain or non-pain task events, which were then convolved with the Glover hemodynamic function with temporal derivatives. Nuisance regressors included: (1) the six motion parameters, their mean-centered squares, their derivatives and their squared derivatives; (2) the mean of cerebrospinal fluid; (3) motion outliers, which were defined as either framewise displacement (FD) > 0.5 mm or std_dvars > 1.5; (4) non-steady outliers from fmriprep. A high-pass filter of 128 s was also applied. Autocorrelation was addressed with an AR(1) model.

In Dataset 3, regressors in the first-level analysis included eight conditions (4 modalities × 2 intensities) convolved with the canonical hemodynamic response function, its temporal derivatives, and six head motion estimates. Moreover, we high-pass filtered images with a cutoff period of 128 s and accounted for temporal autocorrelations using the first-order autoregressive model (AR(1)).

In Dataset 4, the modeling was performed using CANlab tools based on SPM12. For each trial, a boxcar regressor representing the duration of the pain or non-painful stimulation period was specified and convolved with the canonical hemodynamic response function (HRF). Nuisance regressors included: (1) the six motion parameters, their mean-centered squares, their temporal derivatives, and their squared derivatives; (2) the mean of CSF signals; (3) indicator regressors for signal intensity outliers; (4) regressors modeling linear drift across the duration of each run; and (5) regressors modeling task components not of interest, including the cue preceding stimulation, the anticipatory period before pain administration, separate regressors for low- and high-temperature heat stimuli, and the rating period.

#### Second-level analysis

CANlab toolboxes (https://github.com/canlab/CanlabCore) and custom MATLAB (ver R2024b) scripts were used to perform second-level analyses. Single-trial beta maps for each stimulus were averaged for each intensity level and participant and were submitted to one-sample t-tests in Datasets 1 and 2. T-maps for each intensity level were converted to BF maps with Jeffreys-Zellner-Siow priors^113^. To identify regions encoding stimulus intensity and perceptual ratings, we also ran linear regression with single-trial beta maps as regressands and stimulus intensity (linearly coded, e.g., -1, 0, 1 for Dataset 1) or ratings as regressors; the resulting slopes were then submitted to one-sample t-tests. In Dataset 3, beta maps for each stimulus condition were submitted to one-sample t-tests and T-maps for each condition were converted to BF maps with Jeffreys-Zellner-Siow priors. We ran subject-wise analyses in Dataset 4, and thus standard second-level analyses were not applicable. Note that a gray matter mask was applied for all analyses, and that all analyses were conducted in the volumetric space and projected to the surface for visualization purposes only.

#### BF-based voxel classification

Voxel classification was based on BF maps converted from second-level t-maps for Datasets 1, 2, and 3, and first-level t-maps for Dataset 4. The four datasets consisted of varying sample sizes, leading us to choose slightly different BF thresholds for them. Given that the absolute t-values cannot be negative, the conventional cutoff for strong evidence against the null hypothesis, namely 0.1, would require a minimum sample size of 125^114^, which is greater than the sample sizes of Datasets 1 and 2. We thus thresholded log(BF) maps at 0.8 and -0.8 for Datasets 1 and 2, which indicate moderate to strong evidence for or against a statistical effect. In other words, a voxel with log(BF) ≥ 0.8 would be classified as “showing an effect of activation/deactivation compared with the implicit baseline”, whereas a voxel with log(BF) ≤ -0.8 would be regarded as “showing no effect”. For log(BF) = 0.8 and -0. 8, the corresponding t-values are approximately 2.94 and 0.79 when the sample size is 89 (Dataset 1), and 2.96 and 0.92 when the sample size is 11 3 (Dataset 2). Given these sample sizes, t-values of 2.94 and 2.96 yield two-sided p-values of around 0.004, which are smaller than the threshold corresponding to an FDR-corrected significance level of p(FDR) = 0.01, approximately equivalent to an uncorrected p-value of 0.006 in our pain data. In other words, log(BF) = 0.8 is a more stringent threshold than p(FDR) = 0. 01 in Datasets 1 and 2.

To identify systems showing different pain encoding, we classified voxels passing the thresholds into eight classes: (1) “High only+”, showing activation only in the high-intensity conditions (t > 0, log(BF) ≥ 0.8 for High, but log(BF) ≤ -0.8 for Low); (2) “High only-”, showing deactivation only in the high-intensity condition (t < 0, log(BF) ≥ 0.8 for High, but log(BF) ≤ -0.8 for Low); (3) “Low only+”, showing activation only in the low-intensity condition (t > 0, log(BF) ≥ 0.8 for Low, but log(BF) ≤ -0.8 for High); (4) “Low only-”, showing deactivation only in the low-intensity condition (t < 0, log(BF) ≥ 0. 8 for Low, but log(BF) ≤ - 0.8 for High); (5) “Both+”, showing activation in both conditions (t > 0, log(BF) ≥ 0.8 for High and Low); (6) “Both-”, showing deactivation in both conditions (t < 0, log(BF) ≥ 0.8 for High and Low); (7) “Low+ & High-”, showing activation in the low-intensity condition but deactivation in the high-intensity condition (t > 0 for Low, t < 0 for High, log(BF) ≥ 0.8 for High and Low); and (8) “Low- & High+”, showing deactivation in the low-intensity condition but activation in the high-intensity condition (t < 0 for Low, t > 0 for High, log(BF) ≥ 0. 8 for High and Low). Other logically possible classes (e.g., -0.8 < log(BF) < 0.8) were left out as they were not relevant to our purposes. Furthermore, we mainly focused on “High only+” and “Both+” voxels, since the former supports the topographical expansion of sensory representations and the latter helps us understand how newly recruited voxels are related to voxels commonly activated in high- and low-intensity conditions.

The foregoing voxel classifications were based on two intensity levels, that is, (relatively) High and Low. To further explore whether activated regions expand as the intensity increases, we also conducted another version of voxel classification with all three intensity conditions (i.e., 47 °C, 48 °C, and 49 °C) in Dataset 1 . Three voxel classes were defined: (1) “Low & Med & High+”, showing activation in all three intensity conditions (t > 0, log(BF) ≥ 0. 8 for High, Medium, and Low); (2) “Med & High only+”, showing activation only in the medium- and high-intensity conditions (t > 0, log(BF) ≥ 0.8 for High and Medium, but log(BF) ≤ -0.8 for Low); and (3) “High only+”, showing activation only in the high-intensity condition (t > 0, log(BF) ≥ 0.8 for High, but log(BF) ≤ -0.8 for Medium and Low).

Note that since Dataset 3 had a large sample size of 399, we applied a log(BF) threshold of 1.0 or -1.0. In Dataset 4, we performed subject-wise analyses and set the log(BF) threshold to 1.0 or -1.0, given a sufficient number of time points.

#### Permutation test

To test whether the percentage of voxels classified as “High only+” was greater than what could be expected by chance, we performed permutation tests. We randomly permuted intensity labels (Low or High) for all beta maps, and reran second-level analyses and BF-based voxel classification for 1000 iterations to generate a permutation distribution of voxel percentages that were classified as “High only+”.

#### Conjunction analysis

To assess whether “High only+” voxels across datasets and modalities lie in similar regions, we conducted parcel-wise conjunctions. We first parcellated the brain with the CANLab2024 atlas (https://github.com/canlab/Neuroimaging_Pattern_Masks/tree/master/Atlases_and_parcellations/2024_CANLab_atlas), an atlas constructed from 14 previously published atlases, including the Glasser atlas and others^115–134^. We assigned a region to “High only+” if 5% of its voxels were classified as such.

#### Topographical expansion subtyping

To quantitatively examine whether there are different types of expansion, we computed the Euclidean distance between “High only+” voxels and their nearest “Both+” voxels in Datasets 1 and 2. If the distance is small for a certain “High only+” voxel compared with the smoothing kernel, that voxel is likely to have spread from adjacent “Both+” voxels. Otherwise, the voxel is more likely to be recruited anew in response to high-intensity stimulation. We call the first type of expansion “spread expansion”, and the second type “emergent expansion”.

#### Smoothing effect analysis

To assess the effect of spatial smoothing, we re-smoothed Dataset 2, whose unsmoothed data were readily available, with the following kernels: 0 mm (i.e., no smoothing), 2.5 mm, 7.5 mm, and 10 mm. If a voxel is classified as “High only+” even when no spatial smoothing is applied, smoothing should have no substantial effect on it being classified as “High only+”. It is noteworthy that emergent expansion as defined above is unlikely to be an artifact driven by spatial smoothing or vascular mislocalization, as spatial blurring mainly affects voxels close to voxels with true signal.

## Data and code availability

Datasets 1 is available at https://github.com/canlab/canlab_single_trials, and Dataset 2 is available at https://openneuro.org/datasets/ds005256. Dataset 3 is available at https://osf.io/y2n34/ (but referred to as Datasets 1&2). Dataset 4 is from an ongoing study and will be published in the future, but the data needed to replicate findings in the current study, including Dataset 3 and others, are available at https://osf.io/9j7ey. Code for main analyses is also available at https://osf.io/9j7ey.

## Acknowledgments

We thank Dr. Heejung Jung’s and Prof. Hedwig Eisenbarth’s invaluable contribution to data collection and analysis. This work was supported by NIH R01EB026549 (M.A.L. and T.D.W.) and R37MH076136 (T.D.W.).

## Author contributions

Conceptualization: TDW, LBZ; Methodology: LBZ, TDW; Investigation: LBZ, AD, LH, LL; Formal analysis: LBZ, AD, PS, LL; Visualization: LBZ, TDW; Writing – original draft: LBZ, AD; Writing – review & editing: LBZ, AD, LH, PS, LL, ML, TDW; Funding acquisition: TDW, MAL; Project administration: TDW; Supervision: TDW.

## Declaration of generative AI and AI-assisted technologies in the writing process

During the preparation of this work the authors used GPT-5 by OpenAI in order to improve the readability and language of the manuscript. After using this tool, the authors reviewed and edited the content as needed and take full responsibility for the content of the published article.

