## Supplementary material for "Intensity-dependent topographical expansion of sensory representations": Figs S1-14 & Tables S1-3

**Supplemental figures**

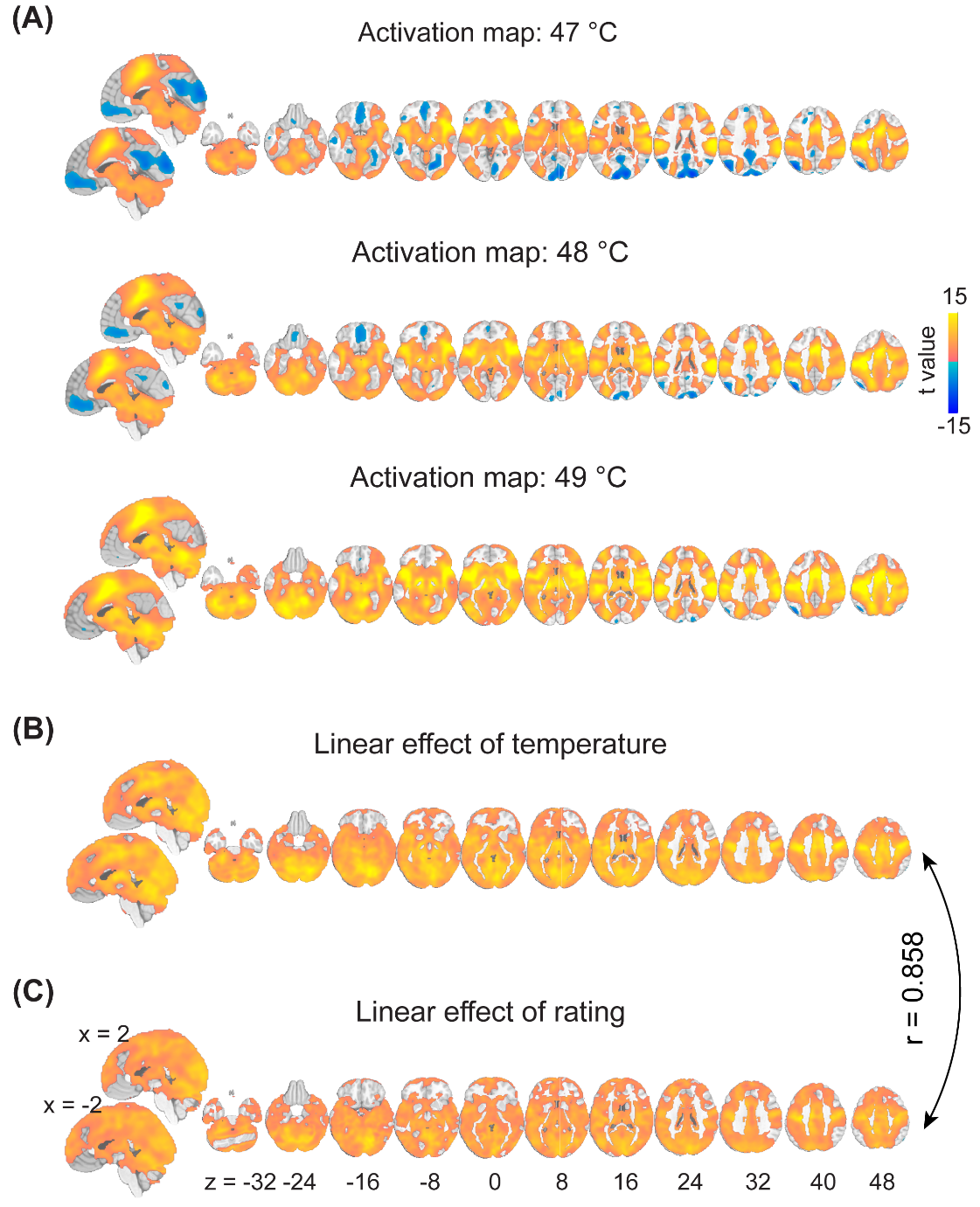

**Figure S1. Heat pain activation in Dataset 1. (A)** Activation maps for three intensity conditions. Heat pain activated a wide range of brain regions. **(B)** Linear effect of temperature. Most activated regions also correlated with stimulus intensity (i.e., temperature). **(C)** Linear effect of ratings. Most brain regions that correlated with stimulus intensity also correlated with pain ratings.

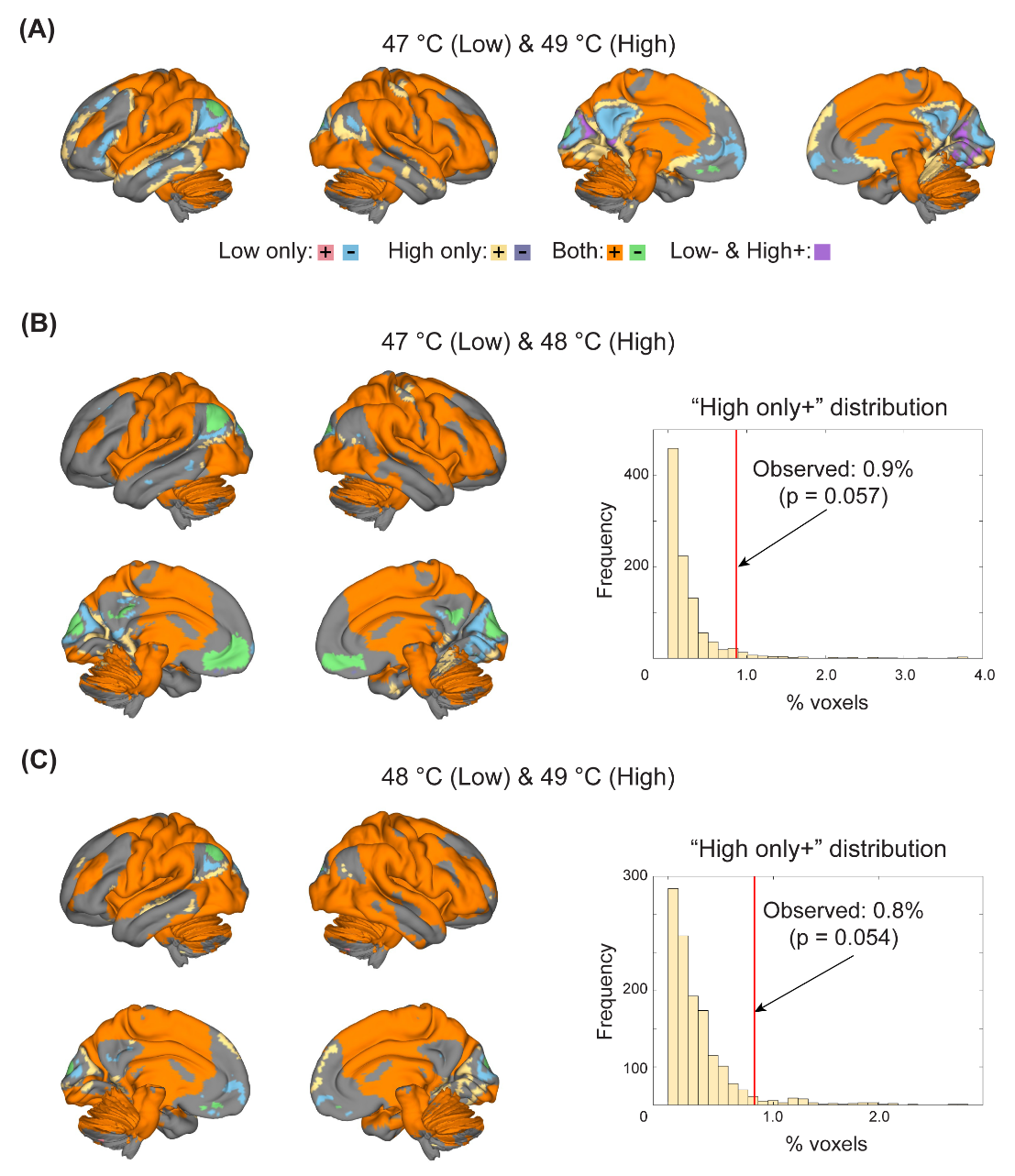

**Figure S2. Voxel classification in Dataset 1. (A)** All classes of voxels in the 47 °C and 49 °C pair. See Figure 2B for the permutation distribution of voxels responding only to the high-intensity condition (i.e., “High only+” voxels). **(B&C)** All classes of voxels in the 47 °C and 48 °C pair (B) and the 48 °C and 49 °C pair (C). The percentages of “High only+” voxels in these two pairs were only marginally greater than the chance level.

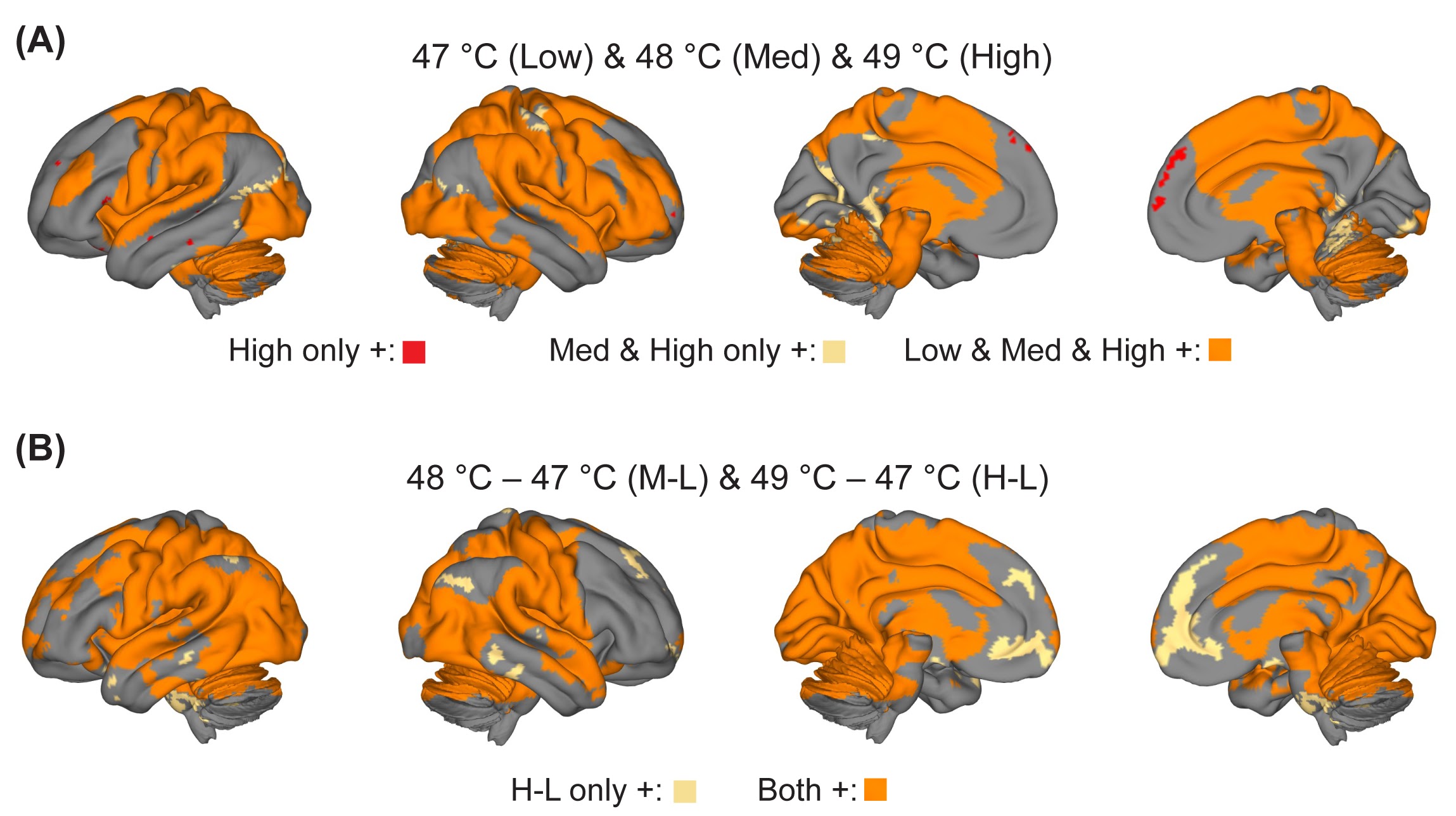

**Figure S3. Topographical expansion across three intensity levels in Dataset 1. (1)** Topographical expansion in all three intensity conditions. While regions such as parts of the S1, cuneus, and cerebellum responded to both the medium- and high-intensity stimuli, a few regions, notably dmPFC, were only activated in the high-intensity condition. **(2)** Topographical expansion using the 47 °C condition as the baseline. Expansion was present in areas such as mPFC.

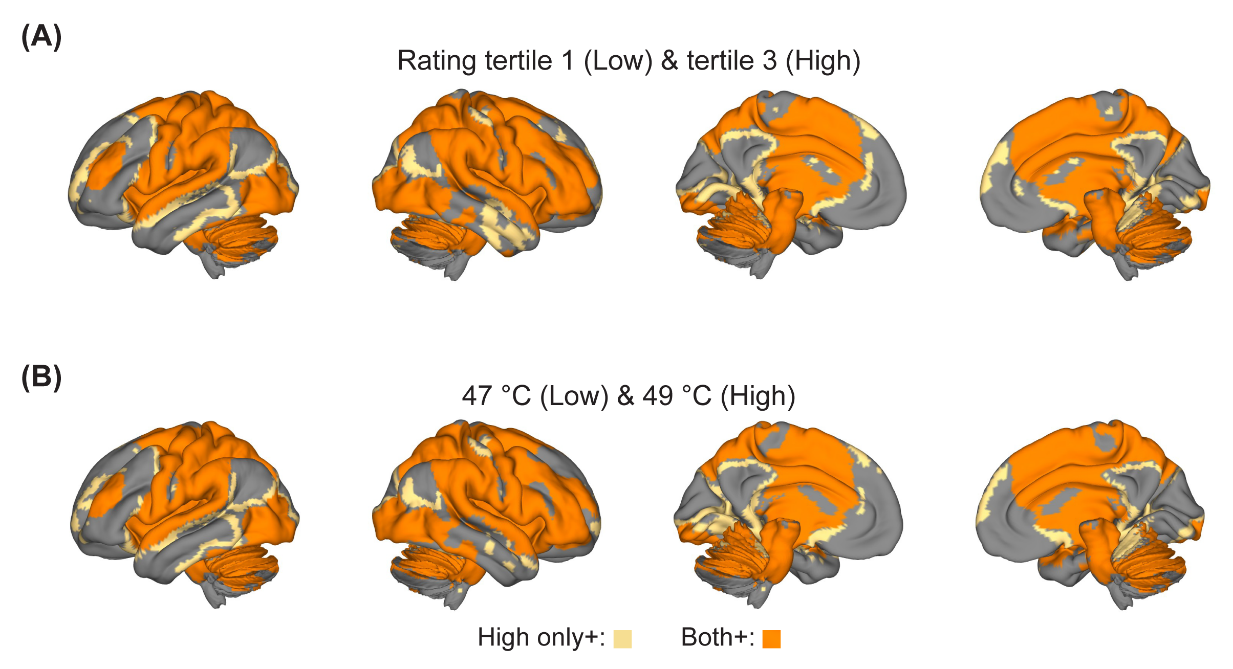

**Figure S4. Rating-based voxel classification in Dataset 1. (A)** Voxel classes in the rating-based 1st and 3rd tertile pair. **(B)** Voxel classes in the intensity-based 47 °C and 49 °C conditions. The distributions of “High only+” and “Both+” voxels were highly similar across these two approaches.

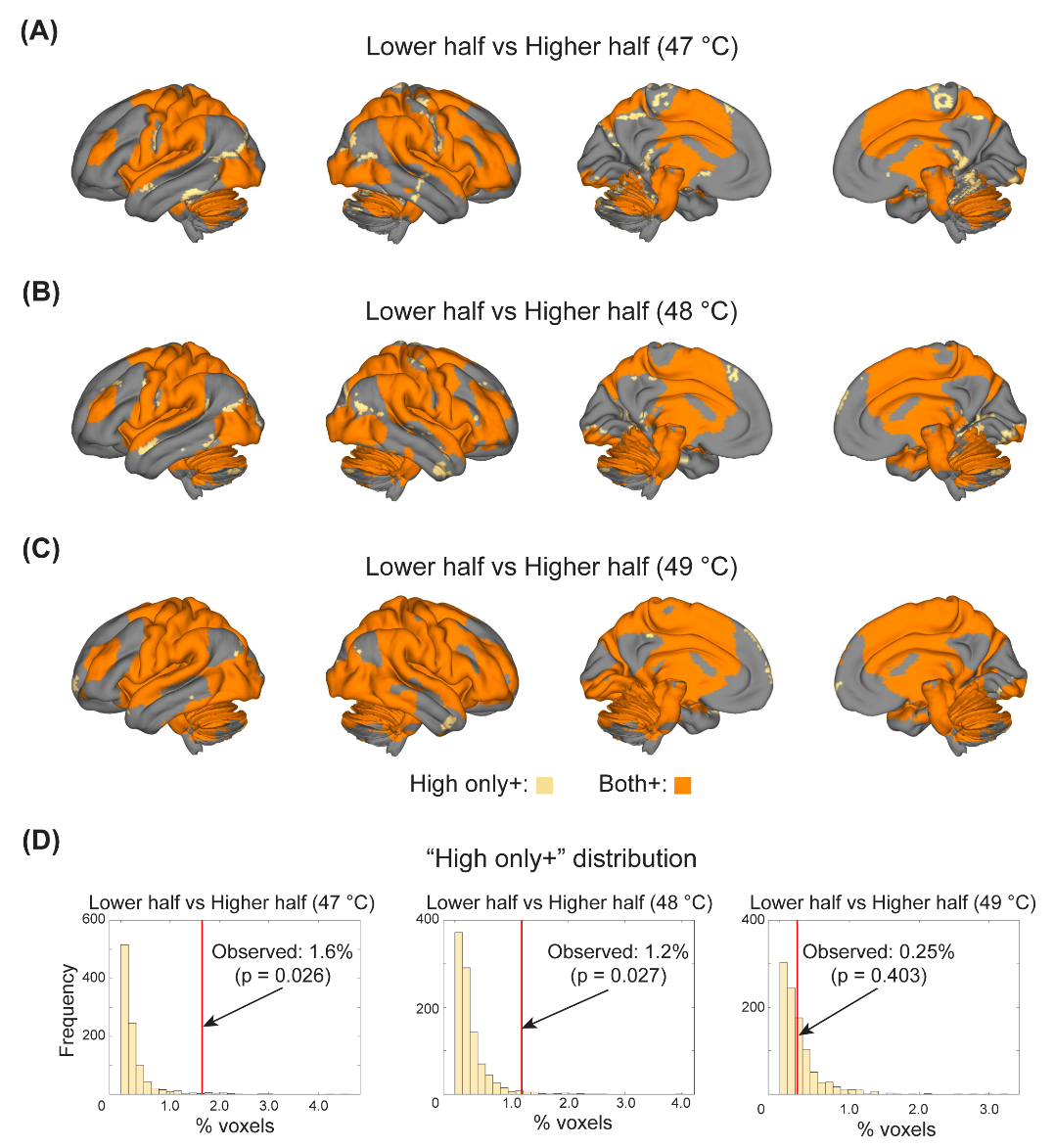

**Figure S5. Rating-based voxel classification while controlling for intensity in Dataset 1. (A~C)** Voxel classes in the rating-based lower- and higher-half trials when stimulus intensity was 47 °C (A), 48 °C (B), and 49 °C (C). **(D)** “High only+” voxels in three temperature-specific pairs. As stimulus intensity increased, fewer “High only+” voxels were found.

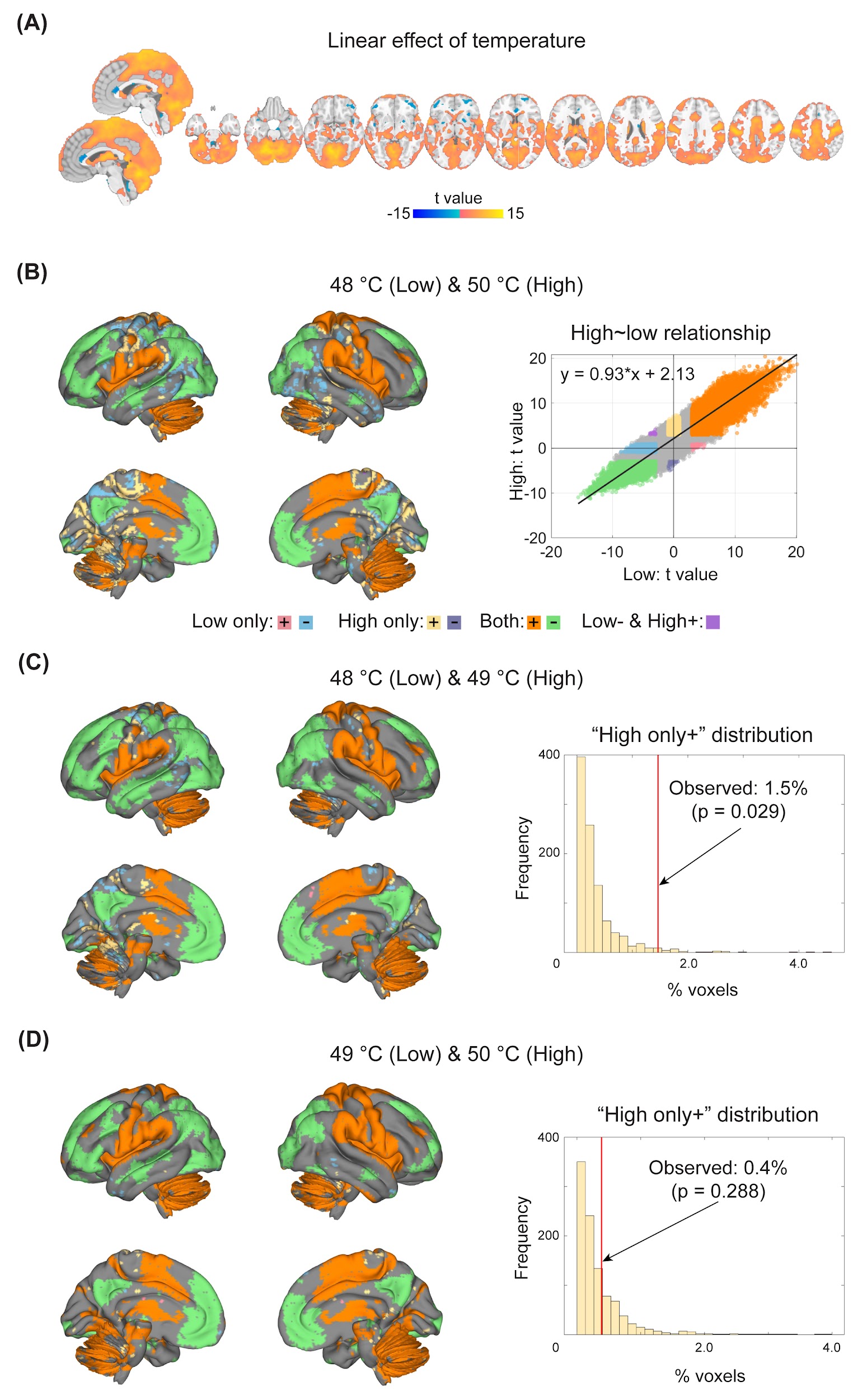

**Figure S6. Voxel classification in Dataset 2. (A)** Linear effect of temperature. Many regions correlated with stimulus intensity (i.e., temperature). **(B)** All classes of voxels in the 48 °C and 50 °C pair. See Figure 3B for the permutation distribution of “High only+” voxels. High-intensity stimuli led to an upward shift in brain activation compared with low-intensity stimuli. Note that the black line in the scatter plot is the fitted line. **(C&D)** All classes of voxels in the 48 °C and 49 °C (C) and 49 °C and 50 °C (D) pairs.

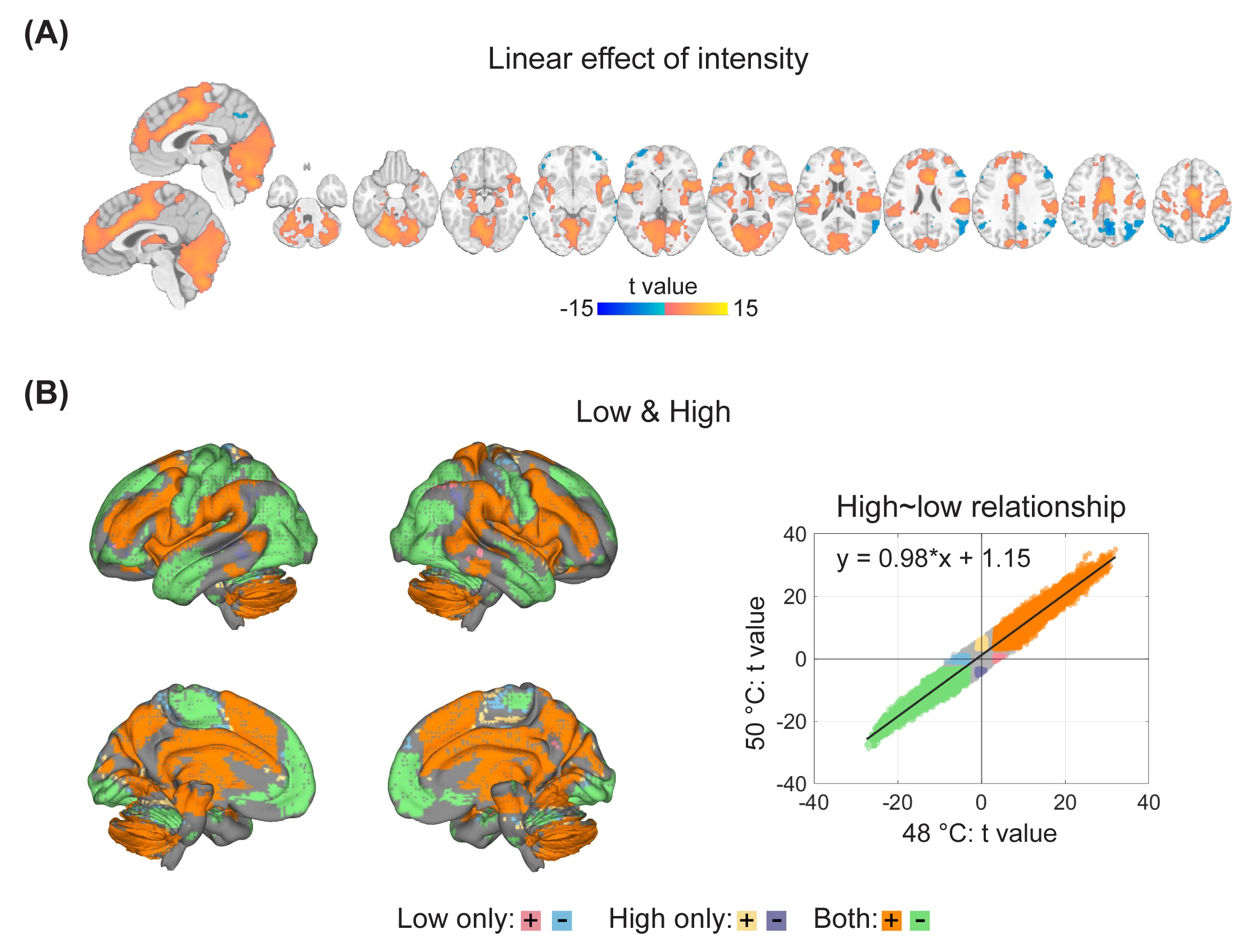

**Figure S7. Voxel classification in Dataset 3. (A)** Linear effect of intensity. Many regions correlated with stimulus intensity. **(B)** All classes of voxels in the low- and high-intensity pair. See Figure 3B for the permutation distribution of “High only+” voxels. High-intensity stimuli led to an upward shift in brain activation compared with low-intensity stimuli. Note that the black line in the scatter plot is the fitted line.

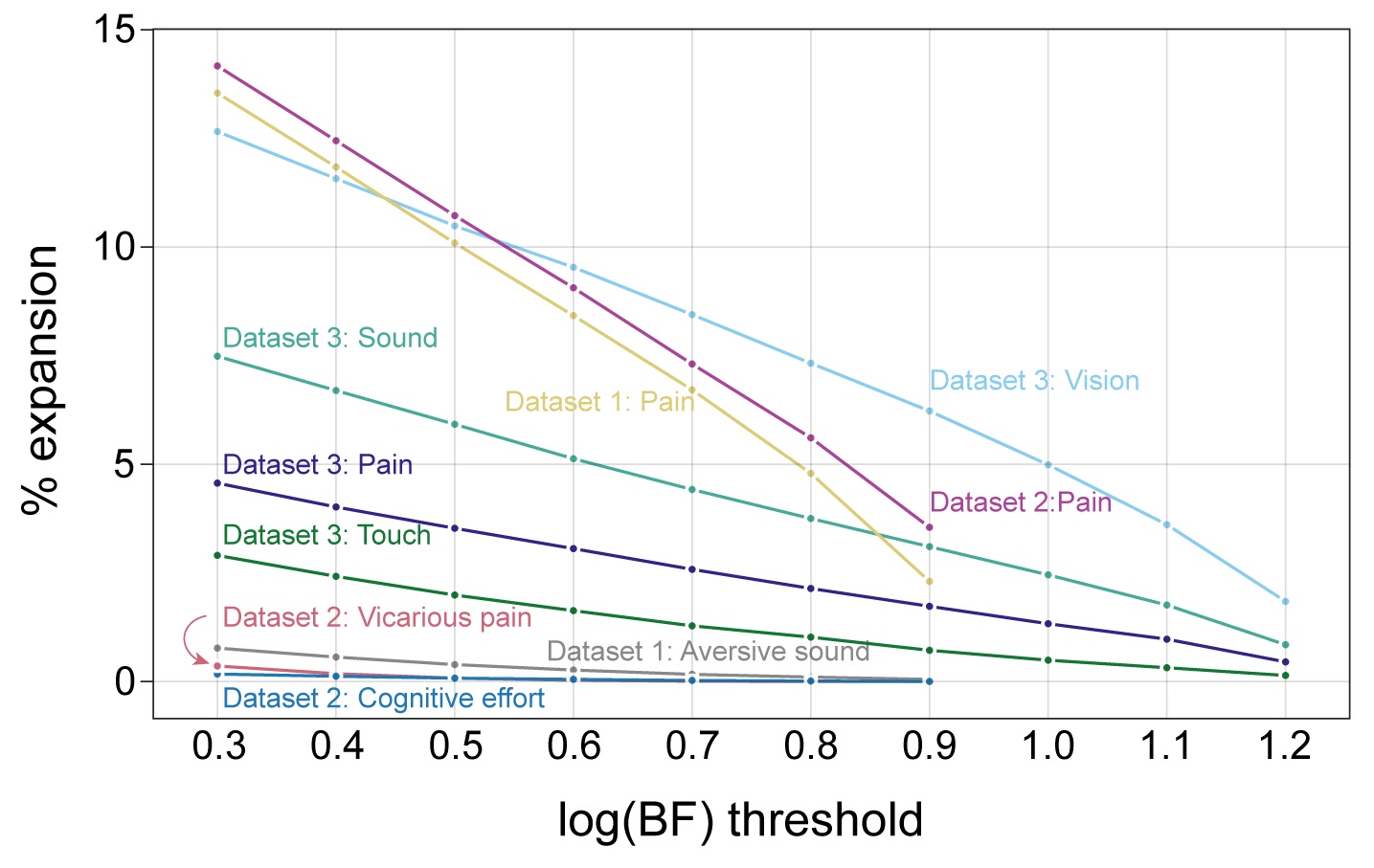

**Figure S8. Sensitivity analysis.** A nearly linear relationship existed between log(BF) thresholds and the extent of expansion. As the threshold loosened, more expansion was observed.

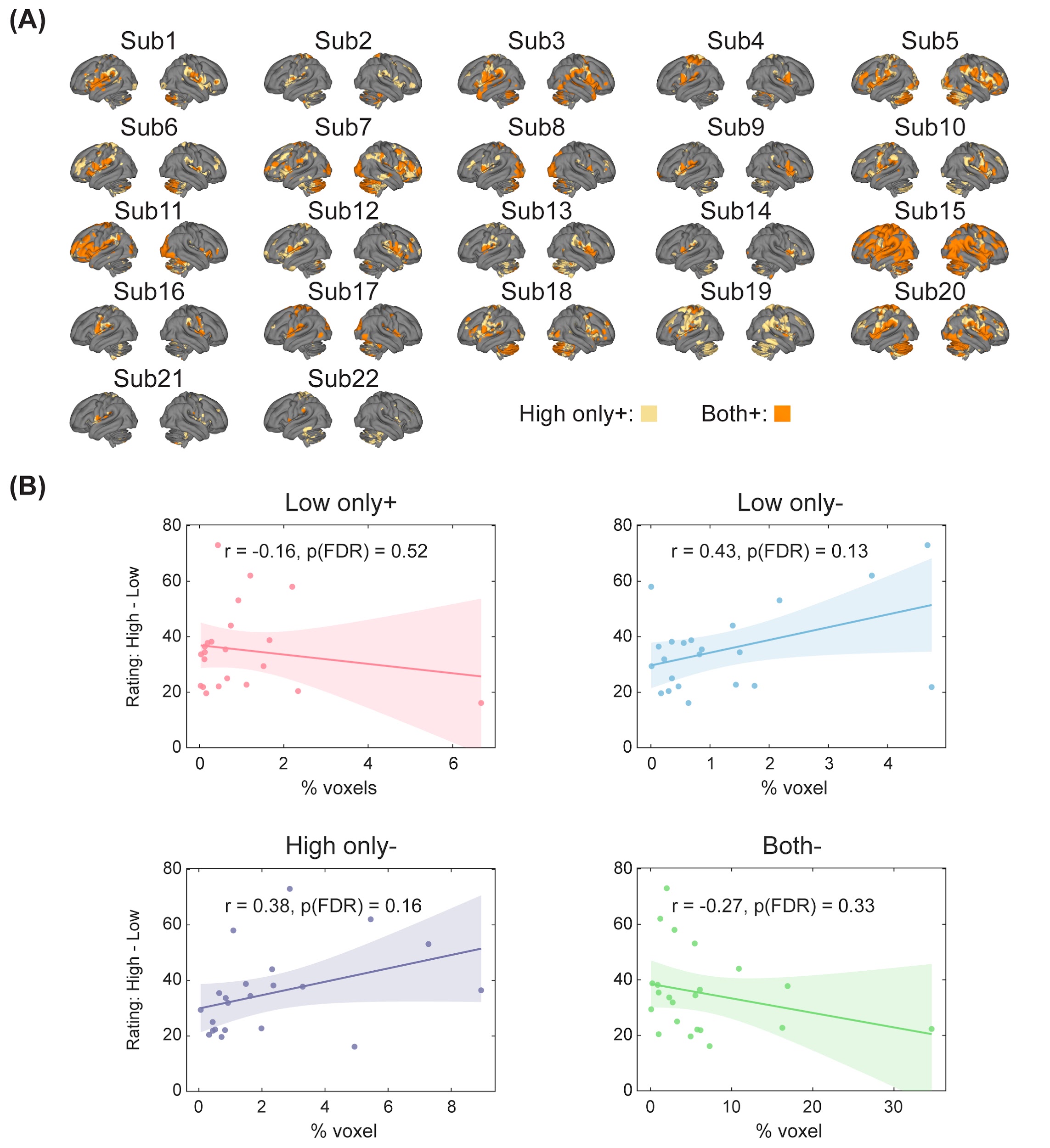

**Figure S9. Individualized expansion maps in Dataset 4. (A)** Voxel classification in all participants. All participants had voxels activated by high-intensity stimuli only, but the precise location and number of voxels differed across individuals. **(B)** Correlation between voxel classes and rating differences. The “Low+ & High-” and “Low- & High+” classes were not analyzed, as a significant number of participants (19 and 13, respectively) had no voxels within these specific classes.
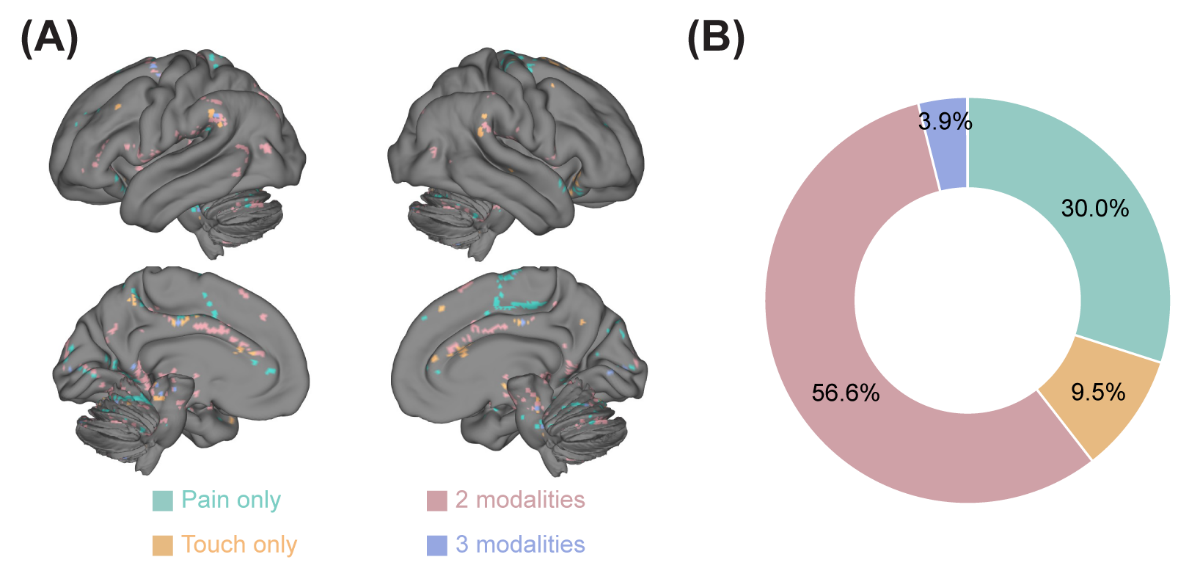

**Figure S10. Voxel-wise conjunction analysis in Dataset 3. (A)** Location of modality-selective and modality-general “High only+” voxels. There were few “High only+” voxels across all four modalities and they were thus not visually discernible in the surface plot. No voxel was selective to audition or vision. **(B)** Percentages of modality-selective and modality-general “High only+” voxels. More than half of the voxels were shared across at least two modalities.

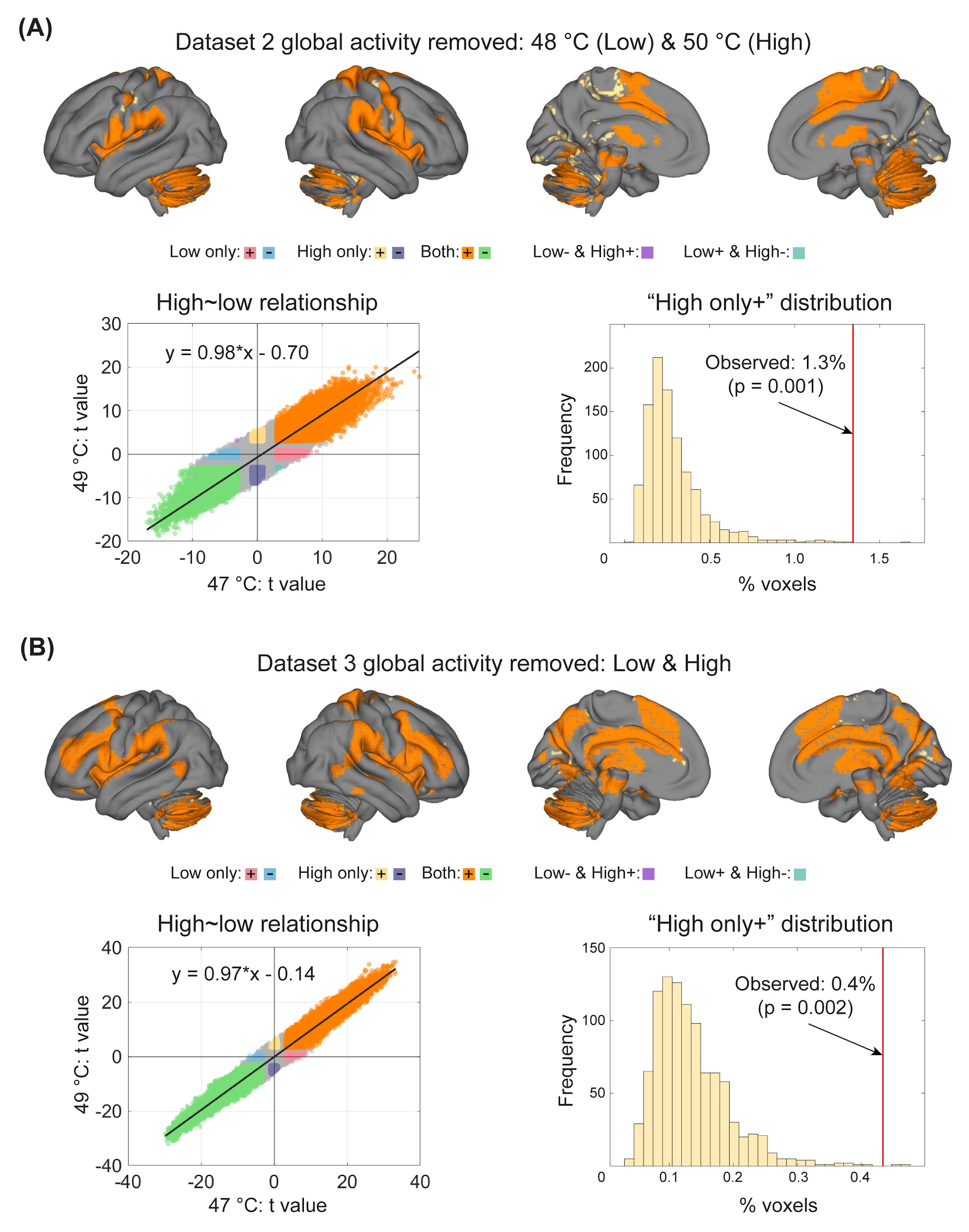

**Figure S11. Pain-induced topographical expansion in Datasets 2 and 3 after global activity removal. (A)** Voxel classification after the removal of global activity in Dataset 2. **(B)** Voxel classification after the removal of global activity in Dataset 3. Note that the black lines in the scatter plots are the fitted lines.

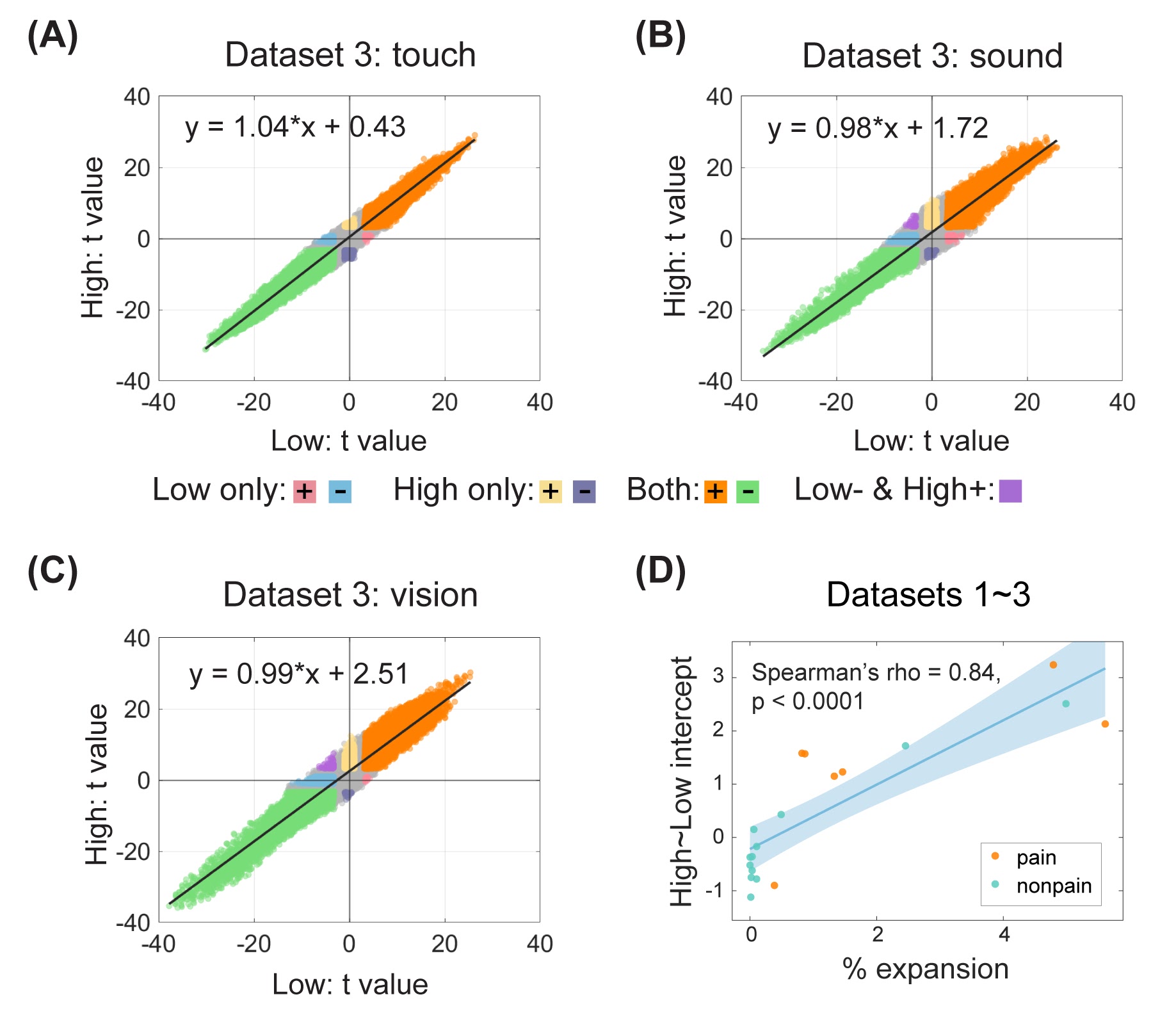

**Figure S12. Global activity shift in non-pain tasks in Dataset 3. (A~C)** Relationship between brain activation in high-intensity stimulation and that in low-intensity stimulation. High-intensity stimuli led to an upward shift in brain activation compared with low-intensity stimuli. Note that the black line in the scatter plot is the fitted line. **(D)** Correlation between topographical expansion and the magnitude of upward shifts of brain activation. Regressing voxel-wise t-values for high-intensity stimuli against t-values for low-intensity stimuli yielded a strong relationship. The intercept from this regression indicates global upwards (for positive values) shifts in activity. Across analyses, contrasts with a higher intercept (global shift estimate) showed more extensive topographical expansion.

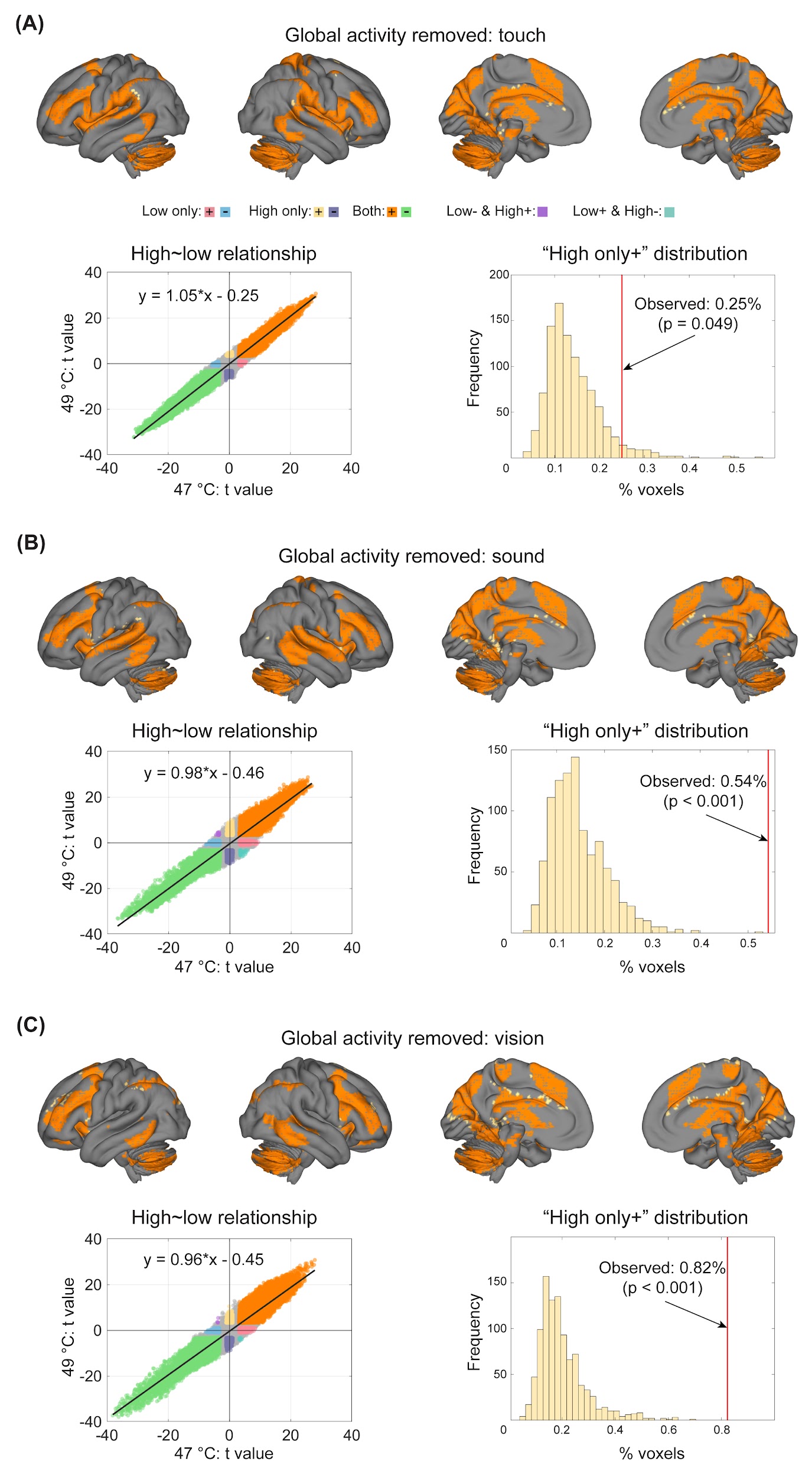

**Figure S13. Non-pain-induced topographical expansion in Dataset 3 after global activity removal. (A~C)** Voxel classification after the removal of global activity for tactile (A), auditory (B), and visual stimulation (C). Note that the black lines in the scatter plots are the fitted lines.

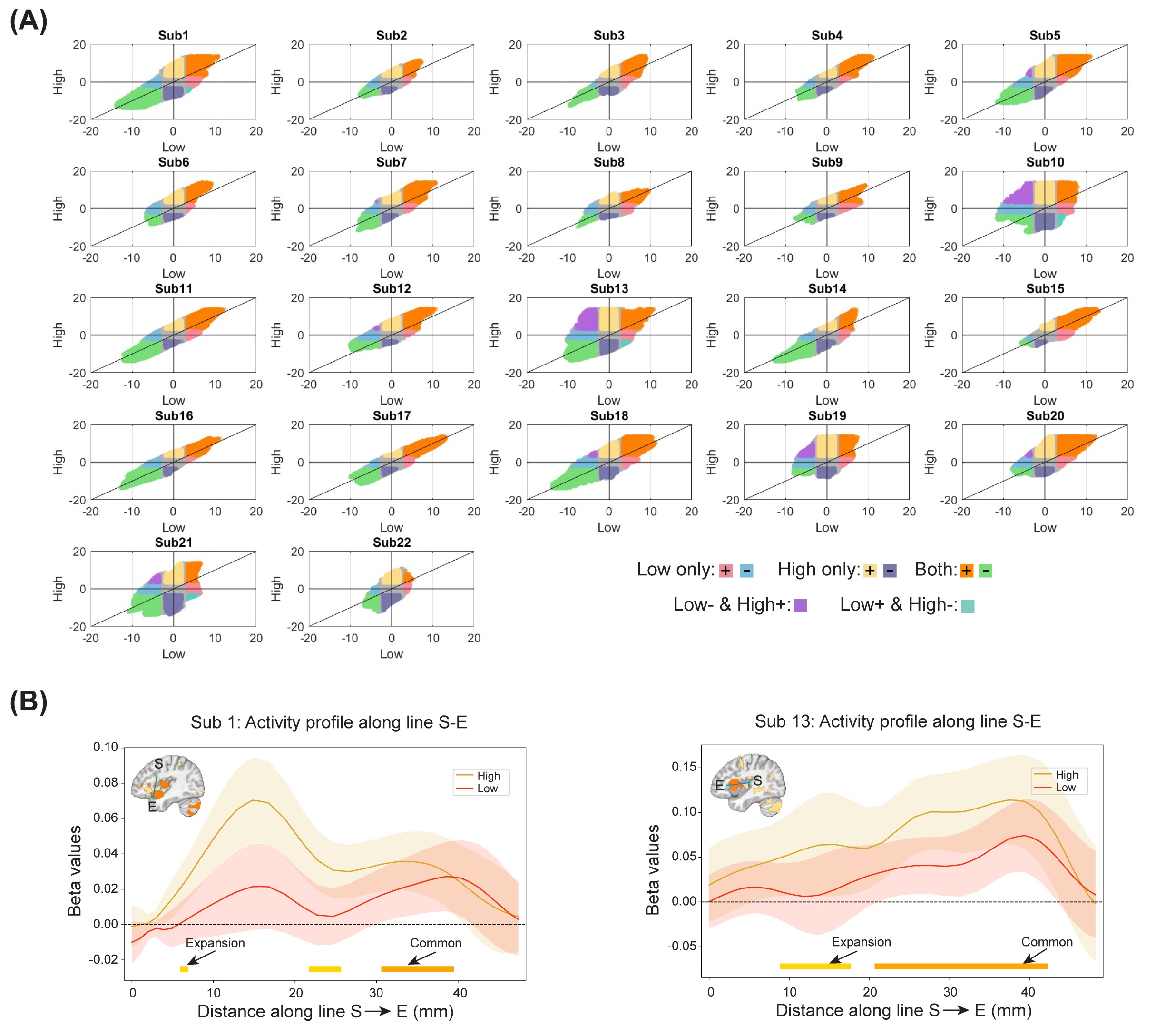

**Figure S14. Individualized topographical expansion in Dataset 4. (A)** Relationship between t-values in the high-intensity condition and low-intensity condition in all participants. Note that the y = x line is plotted, not the fitted lines. **(B)** Example activity profile across “Both+” and “High only+” voxels in two participants. Along the line connecting S and E, brain activity changed systematically across “Both+” and “High only+” voxels. The 99.95% confidence intervals (CI) are plotted because they are close to the α threshold corresponding to log(BF) = 1.0.

**Supplemental tables**

**Table S1. Behavioral ratings in different datasets**

| Datasets | Condition | Mean | SD | Min | Max |
| --- | --- | --- | --- | --- | --- |
| Dataset 1 | Heat pain (L1) | 31.6 | 16.2 | 8.0 | 98.4 |
|  | Heat pain (L2) | 40.6 | 14.8 | 12.6 | 98.5 |
|  | Heat pain (L3) | 48.6 | 16.5 | 14.4 | 100 |
|  | Aversive sound (L1) | 17.7 | 12.6 | 1.3 | 75.1 |
|  | Aversive sound (L2) | 20.8 | 13.8 | 0.2 | 68.7 |
|  | Aversive sound (L3) | 23.7 | 14.9 | 2.4 | 74.0 |
| Dataset 2 | Heat pain (L1) | 51.1 | 28.0 | 8.6 | 142.5 |
|  | Heat pain (L2) | 66.4 | 30.8 | 14.5 | 162.3 |
|  | Heat pain (L3) | 80.0 | 31.4 | 22.2 | 170.1 |
|  | Vicarious pain (L1) | 16.0 | 9.4 | 0.0 | 58.9 |
|  | Vicarious pain (L2) | 22.9 | 11.1 | 0.0 | 72.8 |
|  | Vicarious pain (L3) | 39.9 | 18.8 | 0.0 | 147.0 |
|  | Cognitive effort (L1) | 23.5 | 12.6 | 0.0 | 85.9 |
|  | Cognitive effort (L2) | 31.0 | 15.3 | 0.0 | 105.4 |
|  | Cognitive effort (L3) | 31.9 | 15.0 | 0.0 | 114.0 |
| Dataset 3 | Laser pain (L1) | 3.99 | 1.49 | 0.60 | 8.50 |
|  | Laser pain (L2) | 4.94 | 1.61 | 0.80 | 9.90 |
|  | Touch (L1) | 3.69 | 1.59 | 0.20 | 8.80 |
|  | Touch (L2) | 4.77 | 1.63 | 0.20 | 9.70 |
|  | Sound (L1) | 2.78 | 1.27 | 0.40 | 7.70 |
|  | Sound (L2) | 4.44 | 1.61 | 1.00 | 9.30 |
|  | Vision (L1) | 3.29 | 1.12 | 1.10 | 7.40 |
|  | Vision (L2) | 5.84 | 1.53 | 1.80 | 9.50 |
| Dataset 4 | Heat pain (L1) | 50.9 | 17.6 | 28.7 | 96.1 |
|  | Heat pain (L2) | 86.2 | 20.1 | 56.7 | 118.5 |

**Table S2. Functional MRI acquisition parameters**

| **Parameters** | **Dataset 1*** | **Dataset 1#** | **Dataset 2** | **Dataset 3** | **Dataset 4** |
| --- | --- | --- | --- | --- | --- |
| MRI scanner | Siemens Trio | Siemens Trio | Siemens Prisma | GE MR 750 | Siemens Prisma |
| Magnetic strength | 3T | 3T | 3T | 3T | 3T |
| Field of view (mm) | 248 | 220 | 220 | 192 | 240 |
| Number of slices | 56 | 26 | 56 | 43 | 51 |
| Slice thickness (mm) | 3 | 3.4 | 2.7 | 3 | 2.7 |
| TR (ms) | 460 | 1300 | 460 | 2000 | 1300 |
| TE (ms) | 29 | 25 | 27.2 | 29 | 13.20, 31.45, 49.7 |
| Flip angle (deg) | 44 | 50 | 44 | 90 | 60 |
| Multi-band factor | 8 | / | 8 | / | 3 |
| Slice order | Ascending (interleaved) | Ascending (interleaved) | Ascending (interleaved) | Ascending (interleaved) | Ascending (interleaved) |

* first 25 participants

\# remaining participants

**Table S3. Structural MRI acquisition parameters**

| **Parameters** | **Dataset 1** | **Dataset 2** | **Dataset 3** | **Dataset 4** |
| --- | --- | --- | --- | --- |
| MRI scanner | Siemens Trio | Siemens Prisma | GE MR 750 | Siemens Prisma |
| Magnetic strength | 3T | 3T | 3T | 3T |
| Field of view (mm) | 256 | 256 | 256 | 256 |
| Slice thickness (mm) | 1 | 0.8 | 1 | 0.9 |
| TR (ms) | 2530 | 2000 | 6.896 | 2000 |
| TE (ms) | 1.64 | 2.11 | 2.99 | 2.11 |
| Flip angle (deg) | 7 | 8 | 8 | 8 |
| Slice order | Ascending (interleaved) | Ascending (interleaved) | Ascending (interleaved) | Ascending (interleaved) |
